# Aboral cell types of *Clytia* and coral larvae have shared features and link taurine to the regulation of settlement

**DOI:** 10.1101/2024.10.30.621027

**Authors:** Julia Ramon-Mateu, Anna Ferraioli, Núria Teixidó, Isabelle Domart-Coulon, Evelyn Houliston, Richard R. Copley

## Abstract

Planktonic larvae of many marine invertebrates settle on a suitable substrate and metamorphose into bottom-dwelling adults. Larval settlement is of considerable interest both for ecologists and for evolutionary biologists, who have proposed that anterior sensory systems for substrate selection provided the basis for animal brains. Nevertheless the cellular and molecular regulation of larval settlement, including in Cnidaria (corals, jellyfish, sea anemones, hydroids) is not well understood. We generated and compared anterior (aboral) transcriptomes and single-cell RNA-seq datasets from the planula larvae of three cnidarian species: the hydrozoan jellyfish *Clytia hemisphaerica,* and the scleractinian corals *Astroides calycularis* and *Pocillopora acuta*. Integrating these datasets and characterizing aboral cell types, we defined a common cellular architecture of the planula aboral end, and identified clade-specific specializations in cell types, including unique aboral neural cells in the *Clytia* planula and neurosecretory cell types with distinct molecular signatures in both *Clytia* and coral planulae. Among common planula aboral features were genes implicated in taurine uptake and catabolism expressed in distinct specialized cell types. In functional assays, exogenous taurine inhibited settlement of both *Clytia* and *Astroides* planulae. These findings define a detailed molecular and cellular framework of the planula aboral pole, and implicate localized taurine destruction in defining settlement competence.

## Introduction

Settlement is a key event in the life cycle of cnidarians (jellyfish, hydroids, corals, sea anemones etc; Fig. 1A, B) and other marine species, pivotal in the transition between a pelagic dispersive larval stage and a benthic adult stage. The cnidarian planula larva is morphologically simple, consisting of ciliated ectoderm and endoderm in the form of a tapered cylinder. Cells located at the anterior (aboral) end, the front relative to its swimming direction, are likely to mediate settlement. Aboral but not oral fragments of planulae from the hydroid *Hydractinia* are able to respond to environmental cues and start metamorphosis (Müller et al., 1977). Histological and ultrastructural analyses have described cells at the aboral pole of the cnidarian planula containing secretory vesicles, implicated in substrate adhesion (Thomas et al., 1987). An aboral concentration of putative sensory cells and the presence of an apical nerve plexus have been described in planulae of various species (Attenborough et al., 2019; Nakanishi et al., 2008; Piraino et al., 2011), suggesting that sensory activity and integration of sensory information occur in the aboral region. Planulae of some anthozoan species, including the model *Nematostella vectensis,* bear a presumed sensory organ consisting of apical tuft cells carrying long specialized cilia, surrounded by flask-shaped neural cells (Gilbert et al., 2022; Sabin et al., 2024). In hydrozoan planulae, the broad, aboral end of the larva is enriched with putative neurosensory cells producing GLWamide neuropeptides, known to be active in stimulating settlement and the accompanying drastic cell reorganizations of metamorphosis (Krasovec et al., 2021; Leitz & Lay, 1995; Piraino et al., 2011; Schmich et al., 1998; Takahashi & Takeda, 2015).

**Figure 1.**
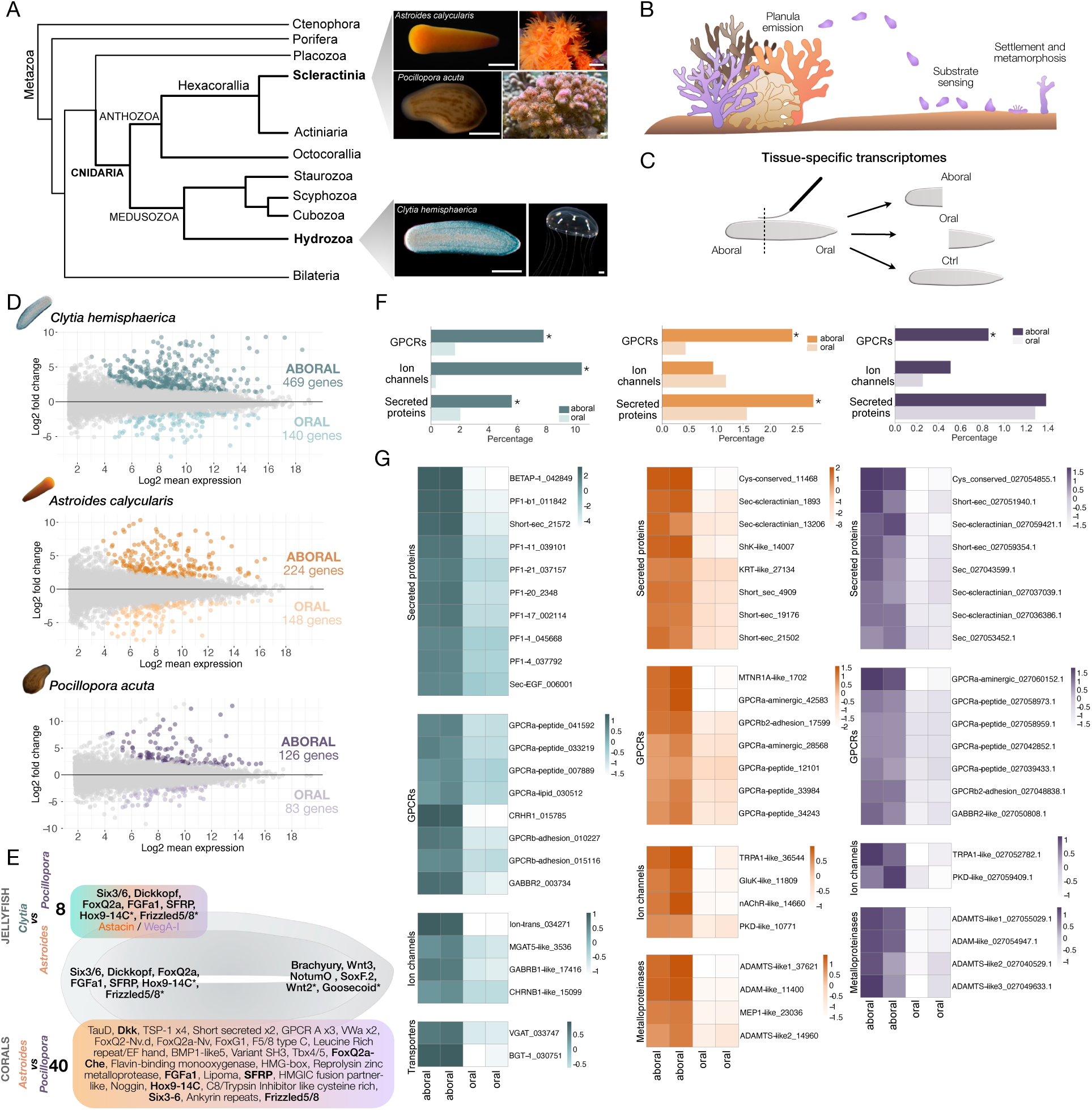
Planula aboral transcriptomes of a hydrozoan and two anthozoan species. (A) Phylogenetic placement of our cnidarian species of study: *Clytia hemisphaerica, Astroides calycularis* and *Pocillopora acuta*. Topology adapted from (Kayal et al., 2018; Schultz et al., 2023). Images of *Clytia* from (Houliston et al., 2010) (reused with permission); the image of the *Astroides* adult colony was taken by Chloé Carbonne. Scale bars, 500 µm for *Astroides* and *Pocillopora* planulae; 100 µm for *Clytia* planula; 1 mm for *Astroides*, *Pocillopora* and *Clytia* adult stages. (B) Schematic illustrating the phases of the settlement response in cnidarians. (C) Schematic of planula bisection. (D) MA plot illustrating the ratio of differential gene expression between aboral and oral ends from each species. Each point on the plot represents a gene, with its x-coordinate indicating the average expression level across all samples and its y-coordinate representing the log fold change in expression between the two conditions. The significantly differentially expressed genes for *Clytia, Astroides* and *Pocillopora* are highlighted in green, orange and purple respectively. Adjusted p-value (padj) < 0.05; FoldChange (FC) > 1. (E) Schematic of a cnidarian planula larva showing shared genes in the aboral-enriched and oral-enriched gene sets obtained from the DGE analysis of planulae of each species. The top panel includes the genes shared between the aboral-enriched sets of *Clytia* and the two corals; the bottom panel includes the shared genes between the aboral-enriched sets of the two coral species. (P-value < 0.05, FoldChange > 1). Asterisks indicate FoldChange > 0 for one or more of the species. (F) Percentage of genes of selected types present in the aboral/oral-enriched gene sets. Asterisks indicate significant differences between the aboral and oral sets (P < 0.05, Fisher’s Exact Test). (G) Heatmaps displaying the gene expression level (normalized counts) in the aboral/oral RNAseq data for selected genes (see full aboral-enriched datasets in Supplementary table 1). Each condition (aboral/oral) includes two replicates. Gene names are based on orthology assignments or Pfam domains when orthology is unclear. Trailing numbers are unique gene identifiers from this project.

A more detailed understanding of the cellular and molecular composition of the planula aboral end would likely yield insights into the settlement response. As well as enabling enquiry into the origins of animal nervous systems which, it has been proposed, may have arisen to coordinate this process (Arendt et al., 2016; Cavalier-Smith, 2017), such knowledge could be valuable in the context of ecosystem management for coral reefs. In this study we generated and compared bulk transcriptomes from the aboral and oral ends of bisected, metamorphosis-competent planula larvae, and single-cell RNA-seq (scRNA-seq) data of the same larval stages. We used three species belonging to the two main clades of Cnidaria: the medusozoan *Clytia hemisphaerica* (jellyfish) and the anthozoans *Astroides calycularis* and *Pocillopora acuta* (brooding stony corals) (Fig. 1A). *Clytia* is a laboratory model species with larvae available from daily spawnings (Lechable et al., 2020). *Astroides* is a mediterranean coral for which larvae were obtained from annual emissions of wild populations, and *Pocillopora* is an indo-pacific, aquarium-propagated tropical coral that emits larvae monthly (Gilis et al., 2014, Massé 2018). We identified and characterized aboral cell types of each species, with a focus on neural and neurosecretory cells. From cross-species comparisons, we identified a conserved aborally expressed gene likely to be involved in taurine catabolism, and we demonstrate that taurine itself inhibits larval settlement. Together, our results allow us to propose a possible mechanism by which taurine mediates settlement competence in cnidarian larvae.

## Results

### Aboral enriched gene sets from the planula of three cnidarian species

We manually bisected settlement-competent planula larvae of *Clytia*, *Astroides* and *Pocillopora* and generated bulk RNAseq data from the separated halves (Fig. 1C; see Methods for details). Principal Component Analysis (PCA) of the most variably expressed genes showed, in all cases, that aboral and oral transcriptomes were distinct (Supplementary Fig. 1). Differential gene expression (DGE) analysis between the aboral and oral ends of each species detected 469, 224 and 126 genes significantly enriched in the aboral transcriptomes of *Clytia*, *Astroides* and *Pocillopora* respectively (Fig. 1D; gene lists provided in Supplementary table 1). The lower number of aborally enriched genes in the coral datasets compared to *Clytia* may, in part, be a consequence of pooling larvae from multiple mother colonies, likely with different genotypes, and larvae at slightly different developmental stages. Fewer genes (140, 148, 83) were detected as preferentially expressed in the oral halves.

### Shared features of aboral transcriptomes across species

Analysis of one-to-one orthologs (see Methods) in common identified a small set of genes that were aborally-enriched across all three species (Fig. 1E). These are all known anterior/apical/aboral expressed genes coding for Wnt and FGF signaling components or transcription factors responsible for patterning the anterior region in cnidarians, protostomes and deuterostomes: the transcription factors Six3/6, FoxQ2A (Hunnekuhl & Akam, 2014; Lowe et al., 2003; Marlow et al., 2014; Santagata et al., 2012; Sinigaglia et al., 2013; Steinmetz et al., 2010; Yankura et al., 2010), and Hox9-14C (Chiori et al., 2009; Gilbert et al., 2024; Marlow et al., 2014; Ryan et al., 2007; Steinworth et al., 2023), the FGFa1 ligand (Gerhart et al., 2005; Rentzsch et al., 2008), a Frizzled 5/8 receptor (Croce et al., 2006; Sinigaglia et al., 2015) and the small secreted antagonists of the Wnt pathway SFRP (Darras et al., 2011; Gilbert et al., 2024; Khadka et al., 2018; Marlow et al., 2014; Pani et al., 2012; Poustka et al., 2007; Sinigaglia et al., 2015) and Dickkopf (Gilbert et al., 2024; Lapébie et al., 2014). These results demonstrate a conserved oral-aboral patterning system across diverse cnidarian planulae and confirm our cutting data are able to accurately identify known aboral markers.

In addition to patterning genes conserved in all three species, we found recurring enriched aboral expression of genes coding for members of other types of protein family. We found genes with potential roles in sensing and signaling as common features of the aboral pole, including a variety of small secreted proteins, G Protein-Coupled Receptors (GPCRs) and ligand-gated ion channels (Fig. 1F).

Among the highest aboral-enriched genes in the *Clytia* planula were several homologous genes coding a family of short secreted proteins, which we term PF1 (Ferraioli et al., in prep.) (Fig. 1G). They do not show sequence homology to any secreted proteins in other species. Analysis of their expression across the *Clytia* life cycle using earlier bulk transcriptome data (Leclère et al., 2019) showed that their expression is almost exclusively at the planula stage, peaking as settlement competency is reached. Distinctly structured secreted proteins of limited taxonomic range were also present in the planula aboral transcriptomes of the two corals.

GPCRs were enriched in the aboral, relative to the oral gene sets in all three species (Fig. 1F). Comparison of the aboral GPCR sets between the three species identified shared receptors between the two coral species but did not find receptors shared between *Clytia* and the corals, either due to the absence of direct one-to-one orthologs or because, when present, these orthologs were not enriched in the aboral region. However, few receptors aboral enriched in *Clytia* did show a corresponding ortholog aboral-enriched in one of the coral species. Notably, both *Clytia* and *Pocillopora* included a metabotropic-like class C GPCR related (although not as a one to one ortholog) to gamma-aminobutyric acid B receptors (GABA_B_R) (Supplementary Fig. 2). Additionally in *Clytia*, we identified a neurotransmitter-gated ion-channel with similarity to GABA_A_ receptors; one amino acid transporter similar to GABA/Glycine vesicular amino acid transporters (vGAT); and one sodium:neurotransmitter symporter channel similar to the sodium-and chloride-dependent betaine/GABA transporter (BGT-1) (Fig. 1G). Sustained activation of GABA_B_R signaling has been found to arrest metamorphosis in the sea anemone *Nematostella* planula (Levy et al., 2021). This suggests a conserved role of GABA (or a related molecule) in the planula nervous system, potentially involved in the regulation of the settlement response in both *Clytia* and the corals.

We noted that all three aboral datasets included genes encoding the Pfam TauD domain (“Taurine catabolism dioxygenase domain”), a subfamily of a broader class of 2-oxoglutarate dependent oxygenases (Islam et al., 2018). The TauD domain is found in enzymes involved in the catabolism of taurine, an amino acid involved in physiological functions, including cell volume regulation, neuroprotection and neurotransmission (Albrecht & Schousboe, 2005; Pasantes-Morales, 1996; Saransaari & Oja, 2000), but also in other proteins. Genes encoding this domain are expressed in the apical organ of *Nematostella* (Sinigaglia et al., 2015) and the apical domain of the planula of two *Acropora* species (Gilbert et al., 2024). Phylogenetic analysis indicates that all are closely related (Supplementary Fig. 3).

### Aboral cells in the *Clytia* planula are specialized neural and neurosecretory types

For scRNAseq we dissociated and fixed cells from *Clytia* planulae as previously described (Ferraioli 2022) at two time points: 52 hpf (late 2-day planulae) and 66 hpf (early 3-day planulae), and processed them using the 10x Genomics platform. We generated four libraries that were merged and integrated with our previous scRNA-seq dataset of the *Clytia* planula larva (Ferraioli 2022) using Harmony (Korsunsky et al., 2019). After filtering steps (see Methods), the final dataset represented 14,627 cells expressing a total of 17,835 genes. Leiden clustering (Traag et al., 2019), generated 27 distinct clusters (Supplementary Fig. 4A, B). These can be grouped into eight broad cell type classes: epidermal, gastrodermal, neural, neurosecretory, interstitial cells (“i-cells”, which are the hydrozoan multipotent stem cells (Bosch, 2009; Siebert et al., 2019), mucous, granular, and cnidocytes (cnidarian stinging cells, developing and mature) (Fig. 2A, B). These are described in more detail elsewhere (Ferraioli 2022 and Ferraioli et al., in prep.). All 27 clusters include cells from the different libraries including the two planula stages (Supplementary Fig. 4C).

**Figure 2.**
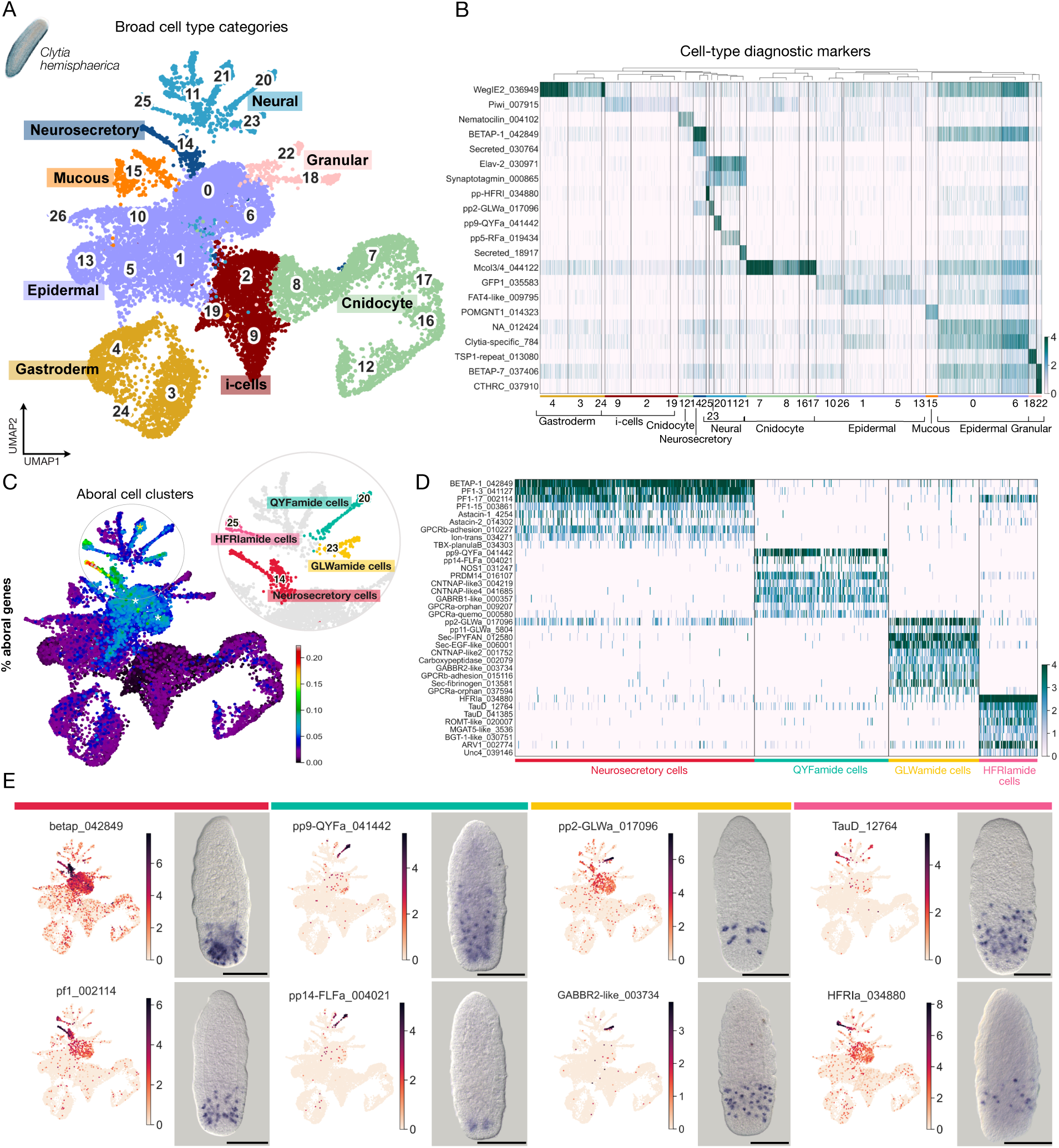
Aboral-enriched cells in the *Clytia* planula. (A) Two-dimensional UMAP representation of *Clytia* planula scRNAseq data with cell clusters color-coded according to eight broad cell type classes: epidermal, gastrodermal, neural, neurosecretory, interstitial cells (i-cells), mucous, granular, and cnidocyte (developing and mature). Codes for class colors are retained in (B). (B) Heatmap of diagnostic marker genes for each cell type. Gene identifiers are plotted on the y axis and cells are plotted on the x axis grouped by the annotation. Gene names are based on orthology assignments or Pfam domains when orthology is unclear. Trailing numbers are unique gene identifiers from this project. See full cluster marker gene lists in Supplementary table 2. (C) UMAP plot of the *Clytia* planula cells, with cells colored based on the percentage of expression of aboral-enriched genes. Warmer colors indicate a higher proportion of aboral gene counts relative to total gene counts per cell. Aboral-gene enriched cell clusters are highlighted in different colors in the upper-right inset. The yellow asterisk indicates the “Nkd2d neural” cluster and white asterisks indicate the two epidermal clusters. P-values for aboral-enrichment significance (Fisher’s Exact Test) are as follows: Neurosecretory cells, 2.47e-72; HFRIamide cells, 8.48e-23; GLWamide cells, 4.12e-20; QYFamide cells, 9.62e-10; Nk2d cells, 8.47e-13; Epidermal cluster 0, 3.04e-10, Epidermal cluster 6, 2.41e-15. (D) Heatmap of marker genes for each aboral cell cluster (complete gene list in Supplementary table 3). Cell-type annotation labels and colors correspond to (C). Gene names are based on orthology assignments or Pfam domains when orthology is unclear. Trailing numbers are unique gene identifiers from this project. (E) UMAP plots highlighting log-normalized expression levels in each cell of selected aboral cell cluster markers, and corresponding in situ hybridization patterns. Scale bars, 100 µm.

To identify aboral cell types, we examined the strength of association between aborally enriched genes (Supplementary table 1) and our cell clusters (Supplementary table 2). Seven cell clusters showed an enrichment in aboral genes (Fig. 2C). Four of these showed a neural signature defined by expression of the pan-neural marker ELAV and other neural-related genes including synaptotagmin-like genes, neuropeptide precursors and ion channels (Cluster 20 = QYFamide cells; Cluster 21 = Nk2d cells; Cluster 23 = GLWamide cells; Cluster 25 = HFRIamide cells). Cells in the Nk2d cluster appear to be heterogeneous. Comparison of gene expression profiles suggests that it may include an aboral subpopulation expressing the Nk2d transcription factor, a short secreted protein, two class A GPCRs and two ion transporters (Supplementary Fig. 5). A fifth cluster was assigned as neurosecretory; it also expressed genes associated with neural function including synaptotagmin-like and calcium signaling related genes but also many secreted proteins.

In addition to these neural/neurosecretory aboral cell clusters, two clusters with epidermal-type signatures (cluster 0 and 6), showed high expression of aboral-enriched genes. A few genes, notably ones coding for short secreted peptides, showed very high expression in these clusters, while others were expressed across many clusters (Supplementary Fig. 6A). In situ hybridization (ISH) for some of these genes revealed expression throughout the aboral epidermis, with more intense signal in a band positioned about two thirds along the oral-aboral axis (Supplementary Fig. 6B). These clusters may thus represent specialized epidermal cells positioned at this location, which are fated to form basal disc epidermis of the primary polyp (Ferraioli et al., in prep.), or precursors of cell types derived from this epidermis. However, the marker gene lists for clusters 0 and 6 also include genes highly expressed in other clusters, so we cannot rule out that they contain cell barcodes associated with ambient mRNA during sample preparation.

### Specializations of the aboral neural and neurosecretory cells

We further investigated the distinct character of each of our *Clytia* aboral cell types. Neurosecretory cells (Cluster 14) were characterized by high expression of many PF1 family secreted proteins, as well as specific expression of two *Clytia*-specific astacin family extracellular metalloproteases (Fig. 2D). ISH revealed dense populations of these cells in the epidermis of the aboral tip of the planula, with a flask-shape morphology and tapered apical section, typical of secretory cells (Fig. 2E). Fluorescent in situ hybridisation (FISH) for a marker gene combined with Hoechst staining (Fig. 3A’) further revealed basally positioned nuclei, in contrast to the apical nuclei of the neighboring epithelial-muscular cells. Basal projections were detected connecting to the apical nerve plexus (Fig. 3A-A’’) suggesting potential signal exchange with other neural cells.

**Figure 3.**
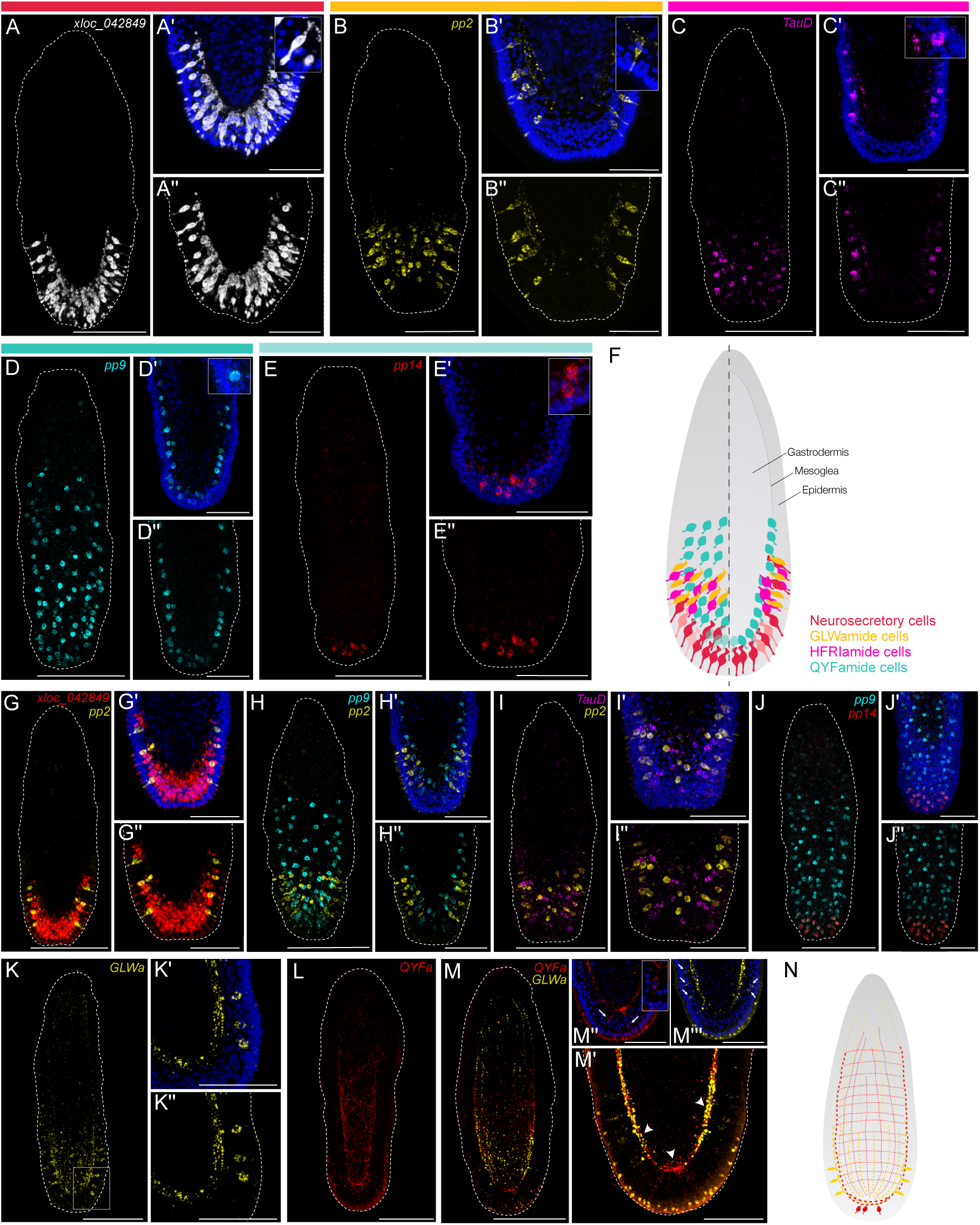
Distinct morphologies of specialized aboral cell types in the *Clytia* planula. (A-B’’) FISH for xloc_042849 (white) and pp2 (yellow). (C-E’’) HCRs for TauD (magenta), pp9 (cyan) and pp14 (red). Magnifications of the aboral ends are shown in (A’-A’’, B’-B’’, C’-C’’, D’-D’’, E’-E’’). Blue staining in (A’, B’, C’, D’, E’) corresponds to Hoechst dye. Insets show close-ups of individual cell bodies. (F) Schematics showing the distribution of aboral cell types in the *Clytia* planula. The left part corresponds to a surface view and the right part corresponds to a medial view. Color code of the cells matches the bars on top of the (A-E’’) panels. (G-J’’) Double HCRs for xloc_042849 (red), pp2 (yellow), pp9 (cyan), TauD (magenta) and pp14 (red). Magnifications of the aboral ends are shown in (G’-G’’, H’-H’’, I’-I’’, J’-J’’). Blue staining in (G’, H’, I’, J’) corresponds to Hoechst dye. (K-M’’’) Neural networks stained with anti-GLWa (yellow) and anti-QYFa (red). (K’, K’’) Magnification of the zone outlined in (K). Note the staining in the apical nerve plexus connecting to individual cell bodies in the epidermis projecting towards the surface. (M’-M’’’) Magnification of the aboral end of (M). (M’’, M’’’) show individual channels combined with Hoechst staining (blue). Inset in (M’’) shows an individual cell body. Arrowheads in (M’) indicate the aboral nerve plexus, while arrows in (M’’ and M’’’) point to individual cell bodies. Note that intense anti-QYFa staining is concentrated at the apical apex and connects to few cell bodies, while the anti-GLWa+ cell bodies are distributed forming a ring at the aboral epidermis below the apical tip. (N) Schematics of the GLWa and QYFa neural networks and cell bodies in the *Clytia* planula. All images are maximum intensity projections of confocal Z-stacks. Images in (B, C, D, E-E’’, H, I-I’’, J-J’’, K) include surface and medial stacks and images in (A-A’’, B’-B’’, C’-C’’, D’-D’’, G-G’’, H’-H’’, K’-K’’, L, M-M’’’) include stacks from only the medial region. Planulae oriented with the oral pole uppermost. Scale bars, 100 µm in full planulae; Scale bars, 50 µm in aboral magnifications.

The GLWamide cells (Cluster 23) express two precursor genes for GLWamide family neuropeptides, pp2 and pp11 (Takeda et al., 2018). GLWamides have strong settlement and/or metamorphosis inducing activity in planulae from several cnidarian species including *Clytia* (Lechable et al., 2020; Takahashi & Hatta, 2011). This cluster also contains cells expressing a gene encoding multiple repeats of the amino acid motif ‘IPYFAN’ followed by dibasic cleavage-like sites, although without the glycine typical of amidated neuropeptides. The GLWamide cells also express five GPCRs, none of which are expressed in other clusters, including a class B GPCR of “adhesion type” and the aboral-enriched metabotropic receptor GABA_B_R-like (Fig. 2D). ISH revealed a ring of sensory-like cells co-expressing pp2 and GABA_B_R-like in the epidermis around the aboral pole (Fig. 2E, Supplementary Fig. 7). These cells have club-shaped bodies with apical processes extending towards the outer surface and basal neurites connecting to the apical nerve plexus (Fig. 3B-B’’). Immunostaining with an anti-LWamide antibody revealed a concentration of neurite processes at the apical end of the larva, with projections extending longitudinally towards the oral end (Fig. 3K-K’’, M).

HFRIamide cells (Cluster 25) are characterized by expression of high levels of a gene which we identify as a putative precursor for a previously undescribed neuropeptide, HFRIamide. The predicted amino acid sequence contains a signal peptide and HFRI sequence followed by a glycine - suggesting amidation (Hayakawa et al., 2019) preceding the dibasic cleavage site. The characteristic protein sequence, including signal peptide and cleavage site, is conserved in other cnidarian species from both Medusozoa and Anthozoa (Supplementary Fig. 8). Other markers for this cluster include the two aboral-enriched TauD genes, as well as the aboral-enriched vGAT-like and BGT-1-like transporters (Fig. 2D). These cells show a similar sensory-like morphology and distribution to the GLWamide cells, forming a ring at the surface of the aboral epidermis (Fig. 3C-C’’). Double HCR, however, shows distinct cell populations (Fig. 3I-I’’).

QYFamide cells (Cluster 20) are defined by the high and specific expression of the neuropeptide precursor gene pp9 (Chari et al., 2021) predicted to generate pyroGluYFamide as well as other amidated neuropeptides. Their transcriptome also shows enriched expression of a nitric oxide (NO) synthase. Additionally, specific markers include two contactin-associated proteins, a zinc finger transcription factor ortholog to the *Nematostella* Prdm14d, shown to be involved in non-ectodermal neurogenesis (Lemaître et al., 2023), and several receptors, including a ligand-gated ion channel with homology to a GABA_A_ receptor subunit, and two class A GPCRs, one of which is very divergent (Fig. 2D). QYFamide cells are located in a zone between the aboral tip and the medial region of the planula. Another neuropeptide precursor, pp14 (likely generating the peptide GPGSRFLFamide) (Chari et al., 2021) shows expression in a subpopulation of the QYFamide cells, concentrated at the aboral tip (Fig. 2E). pp9 expressing cells have a ganglionic-like appearance, distinct from the other neural aboral cell types and characterized by large cell bodies positioned at the base of the epidermis, just above the mesoglea - the basal lamina separating the epidermis from the gastrodermis (Fig. 3D-D’’, E-E’’). Double HCR for pp9 and pp14 confirmed their co-expression in an aboral subpopulation of cells in the neural plexus region (Fig. 3J-J’’). Staining with an anti-pyroGluYFamide antibody showed dense concentration in the aboral nerve plexus and nerve fibers extending the length of the planula body in the mesoglea layer (Fig. 3L). A few individual cell bodies associated with the apical nerve plexus also stained (Fig. 3M’’). Their neurites are organized as an array of perpendicular bundles, in contrast to the GLWamide fibers which run only longitudinally along the planula cell body (Fig. 3M). The observed organization of these ganglionic-type QYFamide cells suggests that they regulate activity of the myofibrils that extend from the basal sides of the ectodermal and gastrodermal epithelial cells in longitudinal and circular directions respectively, to control planula shape.

### Aboral cell types of two coral planula larvae

For *Astroides* and *Pocillopora* we dissociated planula collected from natural emissions and carried out three rounds of 10x Genomics scRNA-seq cell capture for each species. After analysis, we retained 18,964 *Astroides* and 9,733 *Pocillopora* planula cells expressing a total of 21,952 genes and 17,112 genes respectively. Using Leiden clustering, we resolved 31 clusters in the *Astroides* and 30 clusters in the *Pocillopora* planula cell atlases with distinct expression signatures (Supplementary Fig. 9). Clusters were grouped into seven broad cell classes, consistent with previously published cell atlases of anthozoan larvae (*Stylophora pistillata*, (Levy, Elek et al., 2021) and *Nematostella vectensis*, (Sebé-Pedrós et al., 2018; Steger et al., 2022)): epidermal, specialized epidermis, gastrodermal, digestive filaments, neural, secretory, gland mucous, neurosecretory, apical cells and cnidocytes (Fig. 4A, D; Supplementary Fig. 10 and 11).

**Figure 4.**
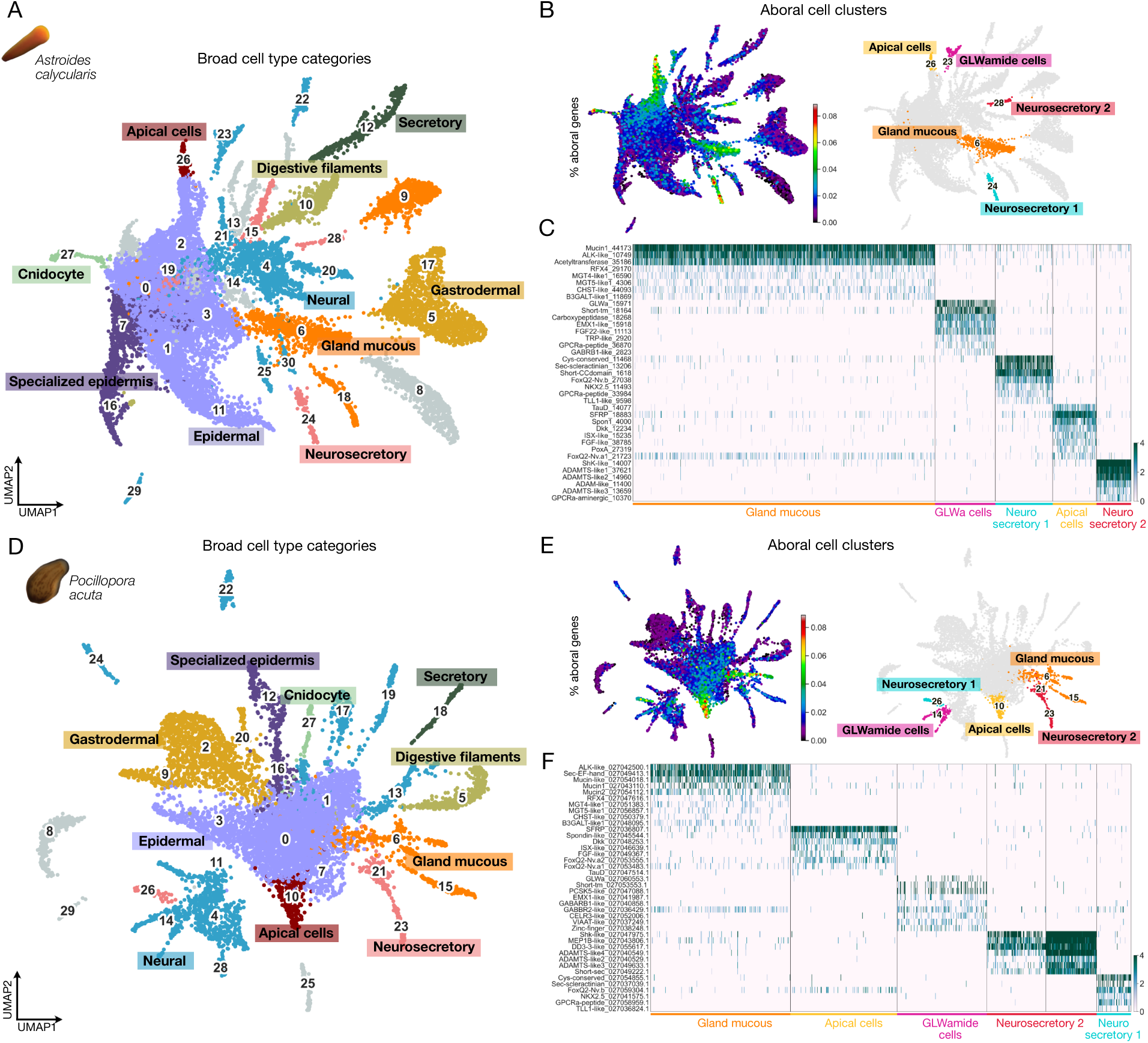
Transcriptomic characterization of aboral cell types in *Astroides* and *Pocillopora* planulae. (A and D) UMAP representation of *Astroides* and *Pocillopora* planula cells respectively labeled by cell type classes: epidermal, specialized epidermis, gastrodermal, digestive filaments, neural, secretory, gland mucous, neurosecretory, apical cells and cnidocytes. Gray clusters have not been annotated. (B and E) Aboral-enriched cell clusters in the *Astroides* and *Pocillopora* planula cell atlas respectively. On the left, UMAP of *Astroides*/*Pocillopora* planula cells, with cells colored based on the percentage of expression of aboral-enriched genes. Warmer colors indicate high proportions of aboral gene counts relative to the total gene counts in each cell. On the right, *Astroides/Pocillopora* planula cell atlas with aboral cell clusters highlighted in distinct colors. P-values for aboral-enrichment significance (Fisher’s Exact Test) were as follows: Gland mucous, 1.75e-06; GLWamide cells 0.71; Neurosecretory 1, 3.38e-22; Neurosecretory 2, 8.97e-05; Apical cells, 1.38e-30 for *Astroides*; Gland mucous_6, 2.71e-05; Gland mucous_15, 0.09; GLWamide cells, 0.73; Neurosecretory 1, 5.06e-05; Neurosecretory 2_21, 6.57e-04; Neurosecretory 2_23, 0.71; Apical cells, 8.92e-20 for *Pocillopora*. (C and F) Heatmap of marker genes for each *Astroides/Pocillopora* aboral cell cluster, with cell-type annotation labels and colors corresponding to (B) and (E) respectively (complete gene list in Supplementary table 3). Gene names are based on orthology assignments or Pfam domains when orthology is unclear. Trailing numbers are unique gene identifiers from this project.

By integrating our *Astroides* and *Pocillopora* aboral transcriptomes with the planula scRNAseq data we identified 5 aboral cell clusters for each species showing equivalent transcriptomic signatures: three cell clusters with a secretory signature (gland mucous, neurosecretory 1 and neurosecretory 2), one neural cluster (GLWamide cells) and one epidermal-like cluster (apical cells) (Fig. 4B, E).

### Transcriptomic and morphological characterization of the coral planula aboral cell types

The *Astroides* and *Pocillopora* planula scRNAseq clustering shows strong similarity, with the *Astroides* dataset containing a larger number of cells. We used the symbiont-free *Astroides* planulae for morphological characterization of the five aboral cell types using *in situ* hybridization chain reaction (HCR) of selected marker genes and antibody staining.

The gland-mucous cell transcriptome is characterized by the expression of mucins and different types of enzymes with potential roles catalyzing post-translational modifications to glycoproteins and glycans including two alpha-1,3-mannosyl-glycoprotein 4-beta-N-acetylglucosaminyltransferase-like proteins, two beta-1,3-galactosyltransferase-like proteins, one alpha-1,6-mannosylglycoprotein 6-beta-N-acetylglucosaminyltransferase-like protein and one carbohydrate sulfotransferase-like protein. These cells also express specifically an ALK-like tyrosine kinase receptor and the RFX4 transcription factor, both of which are also markers for the mucous cells in *Clytia* (Ferraioli et al., in prep.) and the gland mucous cells in the *Nematostella* planula (Steger et al., 2022) (Fig. 4C, F). HCR for a gland mucous cell marker gene shows staining in club-shaped cells densely covering the whole aboral epidermis (Fig. 5A-A’’). They bear apical extensions reaching the surface consistent with a secretory function.

**Figure 5.**
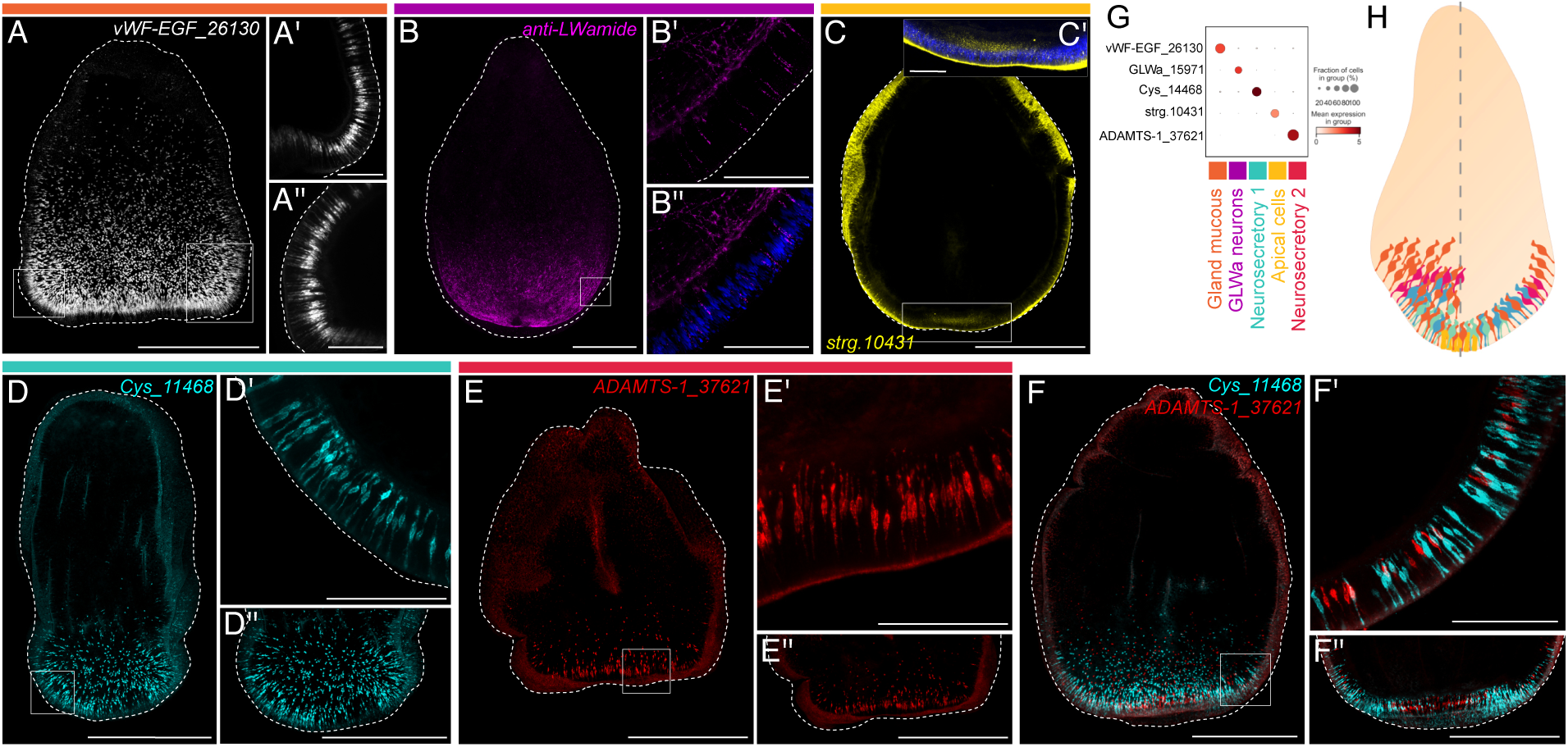
Morphology of aboral cell types in the *Astroides* planula. (A-A’’, C-E’’) HCRs for vWF-EGF_16130 (white), strg.10431 (yellow), Cys_11468 (cyan) and ADAMTS-1_37621 (red). (B-B’’) anti-LWamide staining (magenta). Blue staining in (B’’) and (C’) is from Hoechst dye. (F-F’’) Double HCR for Cys_11468 (cyan) and ADAMTS-1_37621 (red). Squares in (A, B, C, D, E, F) outline the areas shown in higher magnification to the right. All images are maximum intensity projections of confocal Z-stacks. Scale bars, 500 µm in full planulae; 100 µm in the high magnification panels. (G) Dotplot of expression of the marker genes shown in (A-F’’) across the aboral cell clusters. (H) Schematics showing the distribution of aboral cell types in the *Astroides* planula. The left part corresponds to a surface view and the right part corresponds to a medial section. Color code of the cells matches the bars on top of the (A-E’’) panels and the annotation labels in (G).

The two ‘neurosecretory’ clusters show expression of neural markers, including ELAV and the transcription factor AshA (Layden et al., 2012; Steger et al., 2022), and several genes potentially encoding secreted proteins. The “neurosecretory 1” cells are characterized by high expression of two short secreted proteins. One is specific to the Hexacorallia class and, like the PF1 secreted proteins expressed in *Clytia* neurosecretory cells, contains conserved cysteines. The other is exclusively found in other scleractinian coral species (see Supplementary table 3). In both coral species these cells also specifically express a class A GPCRs of peptide-binding type and a metalloprotease related to the Tolloid family, involved in cleaving proteins in the extracellular matrix. They also express two transcription factors: Nkx2.5, identified as apical enriched in *Nematostella* and two *Acropora* species (Gilbert et al., 2024), and a forkhead transcription factor that is a paralog of the aboral FoxQ2a in *Nematostella* (Fig. 4C, F).

The “neurosecretory 2” cells are characterized by the high expression of a cysteine-rich secreted protein with a ShK-like domain, characteristic of the potassium channel-blocking toxin found in the sea anemone *S. helianthus* (Castañeda et al., 1995). These types of domains have also been identified in cnidocyte venom components and in toxin-like neuropeptides in *Nematostella* (Columbus-Shenkar et al., 2018; Moran et al., 2013; Sachkova et al., 2020). This neurosecretory type also shows specific expression of four metalloproteinases of the ADAM family, three of which also detected as significantly enriched in the aboral end (Fig. 4C, F; Fig. 1G). The two neurosecretory cell types show a very similar secretory-like morphology to the gland mucous cells as shown by HCR (Fig. 5D-D’’, E-E’’). They are also concentrated at the apical tip, though in lower numbers. Double HCR for marker genes of each neurosecretory type confirmed expression in distinct populations (Fig. 5F-F’’).

The aboral neural cell cluster is defined by the expression of the GLWamide neuropeptide precursor gene. In both *Astroides* and *Pocillopora*, the transcriptome of these GLWamide cells shows specific expression of neural-related genes, including enzymes involved in the processing of propeptides into their biologically active form; a carboxypeptidase in *Astroides* and a proprotein convertase in *Pocillopora* (Fig. 4C, F). Amongst the few genes that show cluster-specific expression are a neurotransmitter-gated ion-channel with similarity to GABA_A_ receptors (Supplementary Fig. 12), and a homeobox transcription factor EMX-like. Staining with an anti-LWamide antibody revealed a dense nerve plexus at the aboral end of the planula, associated with individual cell bodies within the epidermal layer in an aboral-lateral zone (Fig. 5B-B’’). No cell bodies were detected at the apical tip where neurosecretory cells are more abundant.

The coral planula scRNAseq analysis revealed one aboral cell type clearly distinct from any of those in the *Clytia* planula, the cluster that we term “apical cells”. This cluster expresses the highest proportion of aboral-enriched genes (Fig. 4B, E). Its transcriptomic signature is closely related to the epidermal clusters as shown by the expression of shared marker genes, including a tropomyosin-like and the transcription factor Sox3, expressed in the ectoderm of *Nematostella* (Steger et al., 2022). Among specific apical cell cluster markers are many of the genes forming the gene regulatory network directing the formation of the apical organ in the *Nematostella* planula. These include known aboral/anterior-expressed genes required for the development of the apical organ, including Six3/6, FoxQ2a (Sinigaglia et al., 2013), FGFa1 (Rentzsch et al., 2008) and ISX-like (Gilbert et al., 2022); and other apical markers shown to have specific expression in the apical domain of *Nematostella* planula including Frizzled 5/8, Dickkopf-like, SFRP, Nkx3, Tbx4-5, BMP1-like5 (Gilbert et al., 2024; Sinigaglia et al., 2015). For some of these genes, distinct expression patterns have been observed in one of the two cell types forming the apical organ of *Nematostella* - the apical tuft cells and the larval-specific neurons (Gilbert et al., 2022; Sabin et al., 2024; Sebé-Pedrós et al., 2018). However, all apical organ markers are found co-expressed in the same apical cell cluster of the coral planulae (Supplementary Fig. 13). Comparison to the planula of another scleractinian coral, *Stylophora pistillata,* identified a cluster directly corresponding to the *Astroides* and *Pocillopora* apical cells (Gilbert et al., 2024; Levy, Elek, et al., 2021). Together, these observations indicate homology of the coral apical cells to the *Nematostella* apical organ. A cell type with the “apical cell” signature possibly emerged before the divergence of Actiniaria and Scleractinia, and the subdivision of the apical organ into two distinct cell types evolved subsequently in actinarian sea anemones.

Another notable marker for the coral planula apical cells is a gene containing a TauD domain (Fig. 4C, F), orthologous to the TauD gene identified in the apical domain of *Nematostella* (Sinigaglia et al., 2015) and closely related to the two *Clytia* TauD genes expressed in the HFRIamide cells (Supplementary Fig. 3). HCR for a coral apical cell marker gene in *Astroides* larvae showed signal in a localized region of the epidermis at the most aboral tip (Fig. 5C-C’). The cell bodies are positioned slightly below the aboral surface, their nuclei being less densely packed and positioned deeper than the surrounding epithelial cells (Fig. 5C’). This pattern indicates that specialized epidermal cells at the aboral tip in these coral larvae have a distinct structure.

### Taurine inhibits settlement in both medusozoan and anthozoan planulae

The cnidarian TauD-containing genes emerged from our analyses as a striking common feature of the planula aboral ends, albeit expressed in distinct cell types. The two *Clytia* genes are expressed in the HFRIamide cells, while in the coral larvae, a closely related gene is expressed in apical epidermal-like cells lacking a neural signature. The HFRIamide cells in *Clytia* also express the vGAT-like and BGT-1-like transporters, potentially able to mediate uptake of GABA/Glycine/Taurine related amino acids. The expression of TauD orthologs in specialized cells at the aboral end of other anthozoan planulae (Gilbert et al., 2024; Sinigaglia et al., 2015) suggests a shared and possibly ancestral role across cnidarian larvae. Although the molecular function of these particular TauD proteins remains to be determined, one intriguing possibility is that they are involved in regulating settlement competence via catabolism of taurine. Taurine is an amino-sulfonic acid, widely implicated in many processes of cell metabolism, including osmoregulation, calcium modulation and antioxidation (Huxtable, 1992; Lambert et al., 2015). In the mammalian brain it can act as an agonist at receptors of the GABAergic and glycinergic neurotransmitter systems (Albrecht & Schousboe, 2005), so it is a potential ligand for the GABA_B_R type receptors expressed by in the GLWamide cells. It has been reported to function as a fast neurotransmitter in the scyphozoan “lion’s mane” jellyfish *Cyanea capillata* (Anderson & Trapido-Rosenthal, 2009). Furthermore, studies in hydrozoan larvae have implicated taurine in maintaining the planula state (Müller & Leitz, 2002).

We tested the effect of taurine on larval settlement using *Clytia* in the presence of natural biofilms, and *Astroides* in the absence of active aeration, conditions which both promote settlement. More extensive testing was possible with *Clytia*, for which larvae are readily available from laboratory cultures. Exogenous taurine impaired settlement in both species - with nearly 100% inhibition observed when planulae were incubated in 10^-3^M or 10^-4^M taurine (Fig. 6A, B). Using *Clytia* we could show that the taurine inhibition could be overridden by treatment with GLWamide, placing inhibition by taurine upstream of GLWa secretion (Fig. 6A’). In the hydroid *Hydractinia echinata,* taurine and molecules with N-methyl groups, such as N-methylpicolinic acid (homarine), N-methylnicotinic acid (trigonelline), and N-trimethylglycine (betaine) were detected in high concentrations in the larvae, the levels decreasing upon settlement induction (Berking, 1988), and inhibit metamorphosis when present in micromolar concentrations (Berking, 1986; Walther et al., 1996). Other structurally similar molecules, including GABA and the GABA agonist baclofen, inhibit the planula-to-polyp transition in the sea anemone *Nematostella* (Levy, Brekhman, et al., 2021). In our tests using *Clytia* planulae neither homotaurine, betaine GABA, baclofen or glutamate impaired settlement when present at the same concentrations used for taurine treatments (Fig. 6C), indicating that, at least in *Clytia*, taurine is a specific settlement inhibitor. One possible interpretation of these findings is that specialized aboral cells in cnidarian planulae contribute to regulating settlement by decreasing taurine levels locally, a hypothesis discussed in more detail below.

**Figure 6.**
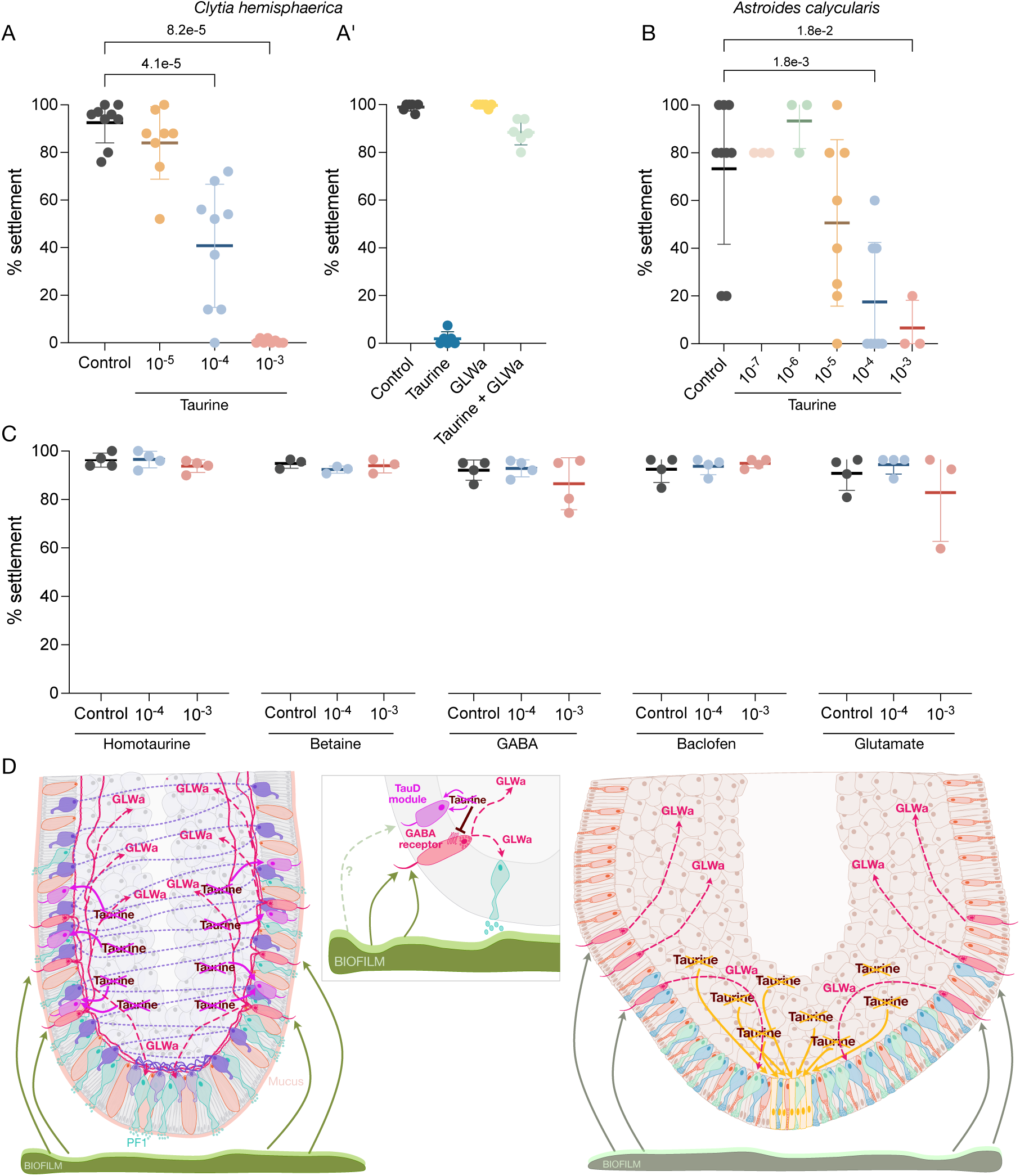
Taurine inhibition of settlement in hydrozoan and coral planulae. (A-A’) Effect of Taurine treatment on *Clytia* settlement. Plots show the means ± standard deviation for each treatment, with concentration units expressed in molarity (M). Each dot represents the percentage of larvae settled in an individual well after overnight treatment, with 100 or 50 planulae used per condition. P-values are indicated for significant differences with respect to the control condition (Mann-Whitney U test). (A’) Combined treatment with Taurine (10⁻³M) and GLWamide (10^⁻6^M). 8 independent experiments were carried out for the Taurine treatment, and 6 independent experiments were carried out for the Taurine + GLWamide combined treatment (see raw treatment data in Supplementary table 4). (B) Effect of Taurine treatment on *Astroides* settlement. Plots show the means ± standard deviation for each treatment, with concentration units expressed in molarity (M). Each dot represents the percentage of larvae that had settled in an individual well on the day when >50% settlement was observed in the control condition, with 5 planulae used per condition. 8 independent experiments were carried out for Taurine at 10⁻⁴M and 10⁻⁵M and 3 for Taurine at 10⁻³M, 10⁻⁶M, 10⁻⁷M (see raw treatment data in Supplementary table 4; see Supplementary Fig. 14 for settlement plots across all days of treatment). P-values are indicated for significant differences with respect to the control condition (Mann-Whitney U test). (C) Effect of other molecule treatments on *Clytia* settlement. Plots show the means ± standard deviation for each treatment, with concentration units expressed in molarity (M). Each dot represents the percentage of larvae settled in an individual well after overnight treatment, with 100 or 50 planulae used per condition. 4 independent experiments were carried out for Homotaurine, GABA, Baclofen and Glutamate treatments, and 3 experiments were conducted for Betaine (see raw treatment data in Supplementary table 4). (D) Proposed model of the cells and molecules involved in cnidarian larval settlement. The schematic depicts the aboral end of the *Clytia* planula on the left and the coral planula on the right. The inset illustrates the proposed regulatory interactions between HFRIamide cells (magenta) and GLWamide cells (red) within the *Clytia* planula.

## Discussion

We have identified shared molecular and cellular features of planula aboral domains of the hydrozoan *Clytia hemisphaerica* and the scleractinian corals *Astroides calycularis* and *Pocillopora acuta*. As illustrated in Fig. 6D and Supplementary Fig. 15, two broad cell-type classes - the secretory-like and the neural-like - have a similar arrangement at the aboral ends of both types of planula.

The secretory cell-types all show flask-shaped morphology with a tapered apical section extending towards the surface to secrete molecules into the environment as previously deduced from histology (Bodo & Bouillon, 1968; Thomas et al., 1987; Vandermeulen, 1974). Neurosecretory cell types in both *Clytia* and coral planulae are characterized by high expression levels of taxon restricted short secreted proteins with similar features: the PF1 secreted proteins in *Clytia,* and a secreted protein with a novel 3D structure in one of the neurosecretory cell types of the coral planula. From the extreme abundance of these proteins, and analogy with other systems such as *Drosophila* (Monier et al., 2024), we speculate that they may have a role in larval adhesion to the substrate. Cnidarian larvae also bear another specialized type of secretory cell, the mucous cells, concentrated in aboral and lateral regions. Mucous secretion could be implicated in slowing down larval swimming at substrate interfaces, promoting adhesion and enhancing bacterial biofilm production to facilitate settlement (Bodo & Bouillon, 1968).

GLWamide cells in both *Clytia* and *Astroides* planulae surround the aboral pole and show the same sensory-like morphology and associated dense neurite network. Enrichment of GLWamide immuno-reactive cell bodies and their neurites has been observed in aboral domains of various medusozoan planulae (Gröger & Schmid, 2001; Leitz & Lay, 1995; Piraino et al., 2011; Schmich et al., 1998) and GLWamide-expressing cells have been detected across an aboral-central domain of the epidermis in the coral *Acropora* planula (Attenborough et al., 2019). GLWamide has been widely implicated in triggering planula cells to undergo metamorphosis, being released downstream of diverse, ill-defined sensory inputs (Müller & Leitz, 2002). The shared expression of genes showing homology to GABA receptors in the GLWamide cells of *Clytia* and coral planulae is thus of note. We identified a GABA_B_R-like in *Clytia* and *Pocillopora* GLWamide cells, and a GABA_A_R-like in both corals *Astroides* and *Pocillopora*. Ectopic activation of presumed GABA_B_ receptors via agonist treatment has been shown to inhibit metamorphosis in *Nematostella* (Levy, Brekhman, et al., 2021). Taken together, these findings suggest a conserved role of GABA-like receptors in the planula nervous system, potentially involved in the regulation of the settlement response. In our assays, GABA itself, unlike taurine, had no effect on settlement in *Clytia*.

Our findings and others have directly implicated taurine in cnidarian planula biology (Berking, 1988; Gilbert et al., 2024; Nakanishi et al., 2008; Sinigaglia et al., 2015). Further, cnidarian larvae for which appropriate data are available show aboral expression of genes encoding Pfam TauD domains (Gilbert et al., 2024; Sinigaglia et al., 2015). These take their name from *E. coli* taurine dioxygenase, although there is no direct evidence of their substrate in cnidarians. Two human genes, γ-butyrobetaine dioxygenase (BBOX1) and trimethyllysine dioxygenase (TMLHE) encode a TauD domain. These are both involved in carnitine biosynthesis, but both have more plausible cnidarian orthologs than the genes we have identified as aborally enriched. In *Clytia*, TauD-expressing cells are neural and co-express genes potentially involved in taurine uptake, including vGAT-like and BGT-1-like transporters. The same TauD encoding genes are also specifically expressed in neurons in the *Clytia* medusa stage (Chari et al., 2021) and it is of note that taurine is an agonist of GABA receptors, close homologs of which we find in GLWamide cells. We have shown that taurine has a common effect in inhibiting settlement in planulae from a coral and *Clytia*. Taking these observations together, we find it reasonable to hypothesize that taurine is a candidate substrate of the TauD proteins.

Putting together the different lines of evidence concerning the aboral TauD protein expression and GLWamide cells, we propose one possible model for settlement regulation (Fig. 6D). High levels of taurine present in larval tissue pre-metamorphosis (Berking, 1988) inhibit spontaneous settlement by binding to metabotropic GABA_B_R-type GPCRs and/or ionotropic GABA_A_R-type receptors in the plasma membranes of GLWamide cells, thereby preventing GLWamide secretion. To allow settlement, aboral cells specialized in the uptake and catabolism of taurine - the HFRIamide cells in *Clytia* and the apical cells in the corals - would act to locally deplete extracellular taurine, thus relieving the inhibition on GLWamide cells and permitting secretion of GLWamide neuropeptides. This mechanism may have been selected during the evolution of planulae to regulate settlement competency, potentially preventing settlement either when the planula is not physiologically ready and/or because environmental conditions are unsuitable. Interestingly, the taurine catabolism genes are expressed in distinct specialized aboral cell types in hydrozoan and coral planulae as well as in the planula of the sea anemone *Nematostella* where it is expressed in the apical organ cells (Sinigaglia et al., 2015). The current situation may have evolved from an ancestral broader expression of taurine catabolism-related genes in an aboral domain, for example, under regulation of aboral domain transcription factors such as Six3/6 or FoxQ2a (Sinigaglia et al., 2013). Throughout the evolution of planulae across diverse species, the TauD proteins may have been selectively retained in different cell types within the aboral end as they diversified independently across species.

This model, in which cnidarian larvae acquire settlement competence via local lifting of taurine mediated inhibitory action around GLWamide secreting cells, integrates our findings but is evidently speculative. Irrespective of the precise mechanisms defining settlement competence, settlement induction is likely to require multimodal input (Birch & Plachetzki, 2023) acting on sensory cells such as GLWamide cells, HFRIamide cells and potentially aboral Nk2d cells. Nitric Oxide (NO) produced by the *Clytia* QYFamide cells is a candidate settlement-inducing signal, these ganglionic-type cells of the aboral neural plexus potentially integrating signals from distinct secreted molecules emitted by the sensory cells. NO is implicated, positively or negatively, in the regulation of settlement and metamorphosis in diverse marine invertebrates (Biggers et al., 2012; Bishop et al., 2008; Bishop & Brandhorst, 2001; Comes et al., 2007; Song et al., 2021; Ueda et al., 2016; Zhang et al., 2012).

Overall this study provides the foundations for a full cellular and molecular understanding of the settlement response in cnidarians through functional analysis of the various candidate receptors and signaling molecules identified in these distinct aboral cell types.

## Methods

### Larva collection

*Clytia* larvae were obtained from fertilizations of laboratory Z strain adults (Leclère et al., 2019). Gametes collected following light-induced spawning of male and female medusae were mixed, and cleaving embryos were transferred into small glass dishes filled with millipore filtered artificial sea water (MFSW-Red sea salt brand) (Lechable et al., 2020). Embryos were left to develop at temperatures of 17-18 °C until the appropriate stage.

*Astroides* larvae were obtained from the release of wild colonies. Fragments of adult female colonies were collected from three sampling sites at the island of Ischia: Sant’Angelo (40°41’31.1’’N, 13°53’35.0’’ E), San Pancrazio (40°42’6.30’’N, 13°57’17.8’’E) and Punta Vico (40°45’33.4’’N, 13°53’1.51’’E), a few days before the full moon of June 2021, 2022 and 2023. They were maintained in a 30 liter tank with water motion provided by a NEWA mini 606 pump and kept in dim-light conditions. Larvae were observed by transparency in the gastrovascular cavity and tentacles of the female colonies (Carbonne et al., 2022). Mature larvae fully developed inside the mother colonies were released 1-2 days after collection. Larvae were transferred into 300 ml plastic bottles filled with seawater and transported to the Villefranche laboratory in less than 24 hours. At the laboratory, larvae were kept in a temperature controlled room at 24°C, in 10 liter plastic containers filled with seawater and constant aeration to prevent settlement. Larvae collected in June 2021 were used for generating the aboral/oral transcriptomic data and the single-cell RNAseq data. Larvae collected in June 2021, 2022 and 2023 were used for immunostaining, in situ hybridizations and molecule treatments. *Pocillopora* larvae were collected from the planulation of parent colonies long-term cultured and asexually propagated by fragmentation at the Oceánopolis Brest aquarium facility (6 tagged colonies, ∼15-20 cm diameter, corresponding to 6 phenotypes of the same genotype). Colonies were placed individually in plankton nets (200 µm mesh) fitted PVC cylinders each evening and transferred back to the holding tank the following morning for a few successive days before the full moon. Night-emitted planula larvae were buoyant and actively swimming, hosting endosymbiotic *Cladocopium sp* dinoflagellates (*Symbiodiniaceae* family) vertically transmitted from each parent colony (Kopp et al., 2016). They were collected with a glass Pasteur pipette, transferred to 300 ml polyethylene bottles filled with 0.2 µm filtered seawater (FSW) (separate batches per each parent colony), and transported back to the laboratory within 2 days from emission (Gilis et al., 2014; Kopp et al., 2016, Massé 2018). FSW was renewed daily (50% of the total volume) and aerated using orbital agitation (Infors incubator, 25 rpm, MCAM laboratory) at 24°C, under a 12-hour day:12-hour night photoperiod (∼70 µmol photons/m²/s, white and blue light) or by gentle air bubbling (LBDV laboratory). Larvae were sorted daily by developmental stage under a stereomicroscope to remove early metamorphic (disk-flattened) stages, retaining only the elongated, swimming planulae. Larvae emitted from individual colonies ‘C3’ and ‘C5’ around the time of the full moon of February 2020 were used for the generation of the aboral/oral bulk RNAseq data. Larvae emitted from ‘C3’ colony around the full moon of December 2022 (6&7/12/2022) and from colony ‘C6’ around the full moon of February 2023 (5&6/02/2023) were used for the generation of single-cell RNAseq data.

### Dissection of aboral/oral ends

Aboral and oral ends of *Clytia* larvae were separated manually using hand-made tungsten wire loops and 2% agarose lined plastic petri dishes. Larvae were anesthetized by placing the dish on ice for 10 minutes, and bisected under a stereomicroscope. For *Astroides* and *Pocillopora*, aboral and oral ends were separated using micro-dissection scissors under a stereomicroscope. Dissected tissues were transferred to the lysis buffer (Ambion, RNAqueous MicroKit), vortexed, immediately frozen in liquid nitrogen and stored at –80 °C until mRNA extraction. *Clytia* larvae were dissected in rounds of maximum 30 minutes, pooled and placed in lysis buffer and snap frozen. *Astroides* and *Pocillopora* larvae were dissected for 5-10 minutes and immediately placed in the lysis buffer and snap frozen.

### Bulk RNA sequencing and differential gene expression analysis

For RNA extraction, tissue samples from several aboral/oral dissections were pooled to ensure sufficient RNA yield for sequencing. For the *Clytia* samples, RNA was extracted from 321 aboral halves, 314 oral halves and 140 uncut larvae for replicate 1, and 320 aboral halves, 317 oral halves and 140 uncut larvae for replicate 2; For the *Astroides* samples, RNA was extracted from 20 oral and aboral halves and 10 uncut larvae for each of the two replicates; For the *Pocillopora* samples, RNA was extracted from 4 aboral halves and 2 oral halves for replicate 1, and 8 aboral halves and 6 oral halves for replicate 2. Total RNA was isolated from each sample using the RNAqueous™ Total RNA Isolation Kit (Ambion, Thermo Fisher). Treatment with DNase I (Q1 DNAse, Promega) for 20 min at 37 °C (2 units per sample) was followed by purification using the RNeasy minElute Cleanup kit (Qiagen). RNA quality of all samples was checked using the Agilent 2100 Bioanalyzer with an Agilent RNA 6000 Nano Kit. Library preparation and sequencing were outsourced (BGI, DNBseq), generating 30 million 100-base paired end reads per replicate. Gene counts for each sample were obtained by mapping the RNAseq reads to the corresponding genomes (a chromosome level genome assembly for *Clytia* (publication in prep.); an in-house generated genome for *Astroides* (see below), and the *Pocillopora damicornis* genome (Cunning et al., 2018) for *Pocillopora*) using STAR (v2.7.11a) (Dobin et al., 2013) with default mapping parameters and the option --quantMode GeneCounts. The gene counts were imported into R and count matrices integrating the gene counts for each condition were generated. Differential gene expression analyses were performed using the DESeq2 R package (Love et al., 2014). Batch effects in *Pocillopora* samples were corrected by adding the batch factor into the design formula. Principal Component Analysis (PCA) and heatmaps were generated using normalized counts obtained with the variance stabilizing transformation (DESeq2 ‘vst’ with blind = F). For the *Pocillopora* dataset, the ‘removeBatchEffect’ function from the limma R package was used to visualize the transformed data with batch variation removed.

### Cell dissociation

For *Clytia*, we prepared single cell suspensions from planulae obtained from two fertilizations using the same batch of mature jellyfish, one let to develop at 17°C until 52 hpf (late 2-day planulae) and the other until 66 hpf (early 3-day planulae). Around 1000 larvae were transferred into a 40 micrometer mesh strainer and placed inside a small plastic petri dish filled with Ca/Mg-free artificial seawater (ASW; 530.5 mM NaCl, 10.7 mM KCl, 3.45 mM NaHCO3, 11.26 mM Na2SO4, pH=8.0). The larvae were washed several times with Ca/Mg-free ASW to remove any residual cations from the seawater. In the final wash, they were transferred to a small petri dish filled with Ca/Mg-free ASW containing 40U/ml ofSUPERase•In™ RNase Inhibitor (Invitrogen), where they were allowed to incubate for 10 minutes. Gentle pipetting with a p1000 tip was used to help dissociate the cells. For the *Astroides* and *Pocillopora* samples, between ten and fifteen larvae were transferred into a 40 micrometer mesh strainer and placed in a small plastic petri dish filled with Ca/Mg-free ASW and 40U/ml SUPERase•In™ RNase Inhibitor (20 U/μL, Invitrogen) for ten minutes. Mechanical disruption of the tissue was achieved by gently rubbing the larvae against the mesh using the interior cap of the plunger of a 1 ml syringe. Once the epidermis was disrupted, the tissue was dissociated by gentle pipetting using a p1000 tip.

In all cases, dissociated cells were recovered in 1 ml of low-Ca /Mg-free ASW (460 mM.NaCl, 9.93 mM KCl, 1.44 mM CaCl2, 10 mM HEPES, pH=7.6) containing 40U/ml SUPERasedn™ RNase Inhibitor. Cell viability following dissociation was evaluated through preliminary testing by diluting the cell suspension with a 1:1 Erythrosin B 0.5mg/ml solution. The dissociation protocol was optimized to ensure a cell mortality rate below 20%. Dissociated cells were immediately fixed in ice cold ACME solution (García-Castro et al., 2021) containing 40U/ml SUPERase•In™ RNase Inhibitor for 30 minutes on ice under rotation. Fixed cells were kept in the ACME solution for up to a month at -20°C until further processing. To wash out the ACME fixative, PBS-1%BSA 0.2 µm filtered containing 40U/ml SUPERasedn™ RNase Inhibitor was added 1:1 to the cell suspension and mixed gently by pipetting. Then, samples were centrifuged at 1000g for 10 minutes at 4°C using a swinging bucket centrifuge. The pellet was resuspended by adding 900 µL of 0.2 µm filtered PBS -1%BSA containing 100U/ml SUPERasedn™ RNase Inhibitor. To cryopreserve cells, 100 μL of DMSO were added per tube and samples were stored at -80°C until performing the single-cell sequencing.

### Single-cell RNA sequencing

Single-cell RNA sequencing (scRNA-seq) for all samples was performed using 10X Genomics technology. Single-cell suspensions kept at -80°C were thawed on ice. To wash out the DMSO, cells were centrifuged twice at 1000g for 10 minutes at 4°C using a swinging bucket centrifuge. The supernatant was discarded and the pellet was resuspended in PBS-1%BSA 0.2 micron filtered containing 100U/ml SUPERasedn™ RNase Inhibitor (20 U/µL) after the first centrifugation and in PBS-0.1%BSA 0.2 micron filtered containing 100U/ml SUPERase•In™ RNase Inhibitor (20 U/μL) after the second centrifugation. Cell concentration was roughly assessed using the Countess 3 Automated Cell Counter (Invitrogen). For two out of the three *Astroides* samples, cells were stained with 1/300 DRAQ5 (Thermo Fisher, stock 5 mM) and sorted using a BD Influx cell sorter to discard debris and dublets. The rest of the *Astroides* samples and the *Clytia* and the *Pocillopora* samples were processed directly after having optically examined the cell suspensions and discarded the presence of debris and cell clumps. Cells were loaded into the 10X Genomics platform for encapsulation with the capture goal of between 7000-10,000 cells per sample. We conducted 4 capture experiments for the *Clytia* samples and 3 captures for the *Astroides* and *Pocillopora* samples. cDNA libraries were prepared according to the Chromium Next GEM Single Cell 3’ library preparation protocol v3.1. Sequencing was performed on an Illumina NextSeq2000 device with NextSeq 2000 P3 Reagents (100 Cycles). Paired end reads length: read1=28bp, Index1=10bp, Index2=10bp and read2=90bp (Index Kit TT SetA).

### Single-cell transcriptomic analysis

Individual 10x sample libraries were demultiplexed using Cell Ranger Makefastq v7.0.0 with default settings. Reads for each sample were mapped using STARsolo (Kaminow et al., 2021) with the --soloCellFilter EmptyDrops_CR option and applying a 300 UMI cutoff using the genomes and gene models described above. Output count matrices were analyzed with Scanpy 1.9.1 (Wolf et al., 2018). To exclude low-quality cells from downstream analysis, we filtered cells with fewer than 200 genes expressed and genes detected in fewer than 3 cells. Additionally, we removed cells with high mitochondrial counts by setting thresholds for filtering based on visual examination of the percentage of mitochondrial counts across cells within each species’ dataset. The percentage of mitochondrial counts was set to a maximum of 20% for *Clytia*, 0.4% for *Astroides* and 5% for *Pocillopora*. Following this initial filtering, we ran each dataset through a standard Scanpy analysis, as follows, using default parameters. Counts per cell were normalized, log-transformed and highly variable genes were computed. Then, datasets were scaled and Principal Components (PCs) were computed on highly variable genes. Clustering was performed by first running the neighbors function using 50 neighbors and adjusting the number of principal components (PCs) individually for each dataset (22 PCs for *Clytia*; 26 PCs for *Astroides*; 23 PCs for *Pocillopora*), then applying the Leiden graph-clustering algorithm (resolution=1 for *Clytia* and *Pocillopora;* resolution=0.9 *for Astroides*) (Traag et al., 2019). Cluster “marker genes”’ were identified by extracting significantly differentially expressed genes per cluster using the Scanpy’s rank_genes_groups function with the Wilcoxon rank-sum test method.

For the *Clytia* analysis, the Harmony algorithm (Korsunsky et al., 2019) was used to correct the batch effects during the integration of datasets generated in this study with previously produced datasets for *Clytia* planula (Ferraioli et al., in prep.).

For *Clytia* and *Astroides*, a cluster with no unique significantly represented genes when compared to other clusters and expression of marker genes across all clusters was removed from the datasets. The resulting datasets were re-clustered and used for all downstream analysis.

### Astroides calycularis genome

Genomic DNA was extracted from the tissue of an isolated polyp of an adult colony. An individual polyp was separated from the colony by breaking its calcareous skeleton using pliers. The isolated polyp was snap frozen in liquid nitrogen and immediately thawed. Polyp tissue was then separated from the skeleton using an airbrush, collected in a tube, snap-frozen in liquid nitrogen, and stored at -80°C until processed for DNA extraction. For DNA extraction, frozen polyp tissue was thawed on ice. 2 mL of DNA extraction buffer (200 mM Tris-HCl pH 8.0, 20 mM EDTA, 0.1% SDS, 0.5 mg ml-1 proteinase K (Qiagen), 0.2 mg ml-1 RNase A (Qiagen) were added and incubated at 50°C for 2 hours until the solution became uniform and clear. An equal volume of phenol was added, mixed by inversion and centrifuged for 30 min at 14,000 rpm (room temperature). Then, the upper layer was transferred to a new tube and the same volume of chloroform was added. The solution was mixed by inversion and centrifuged for 30 min at 14,000 rpm (room temperature). A second chloroform wash was performed. The upper layer was transferred into a new tube and X1/10 volume of 5 M NaCl and 0.7 volumes of isopropanol were added. The solution was gently mixed by pipetting, incubated for 30 minutes at room temperature and centrifuged for 40 min at 14,000rpm (room temperature). The DNA precipitate was rinsed with ice cold 70% ethanol, dried and dissolved into TE buffer (10 mM TrisHCl pH 8.0, 1 mM EDTA pH 8.0). A total of 30 μg of DNA were obtained from 28 mg of polyp tissue. In collaboration with Marie-Jeanne Arguel at the France Genomix platform, IPMC, Sophia Antipolis, longer DNA fragments were selected using The Short Reads Eliminator (SRE) Medium kit (Circulomics). Sequencing library preparation was performed at this platform using the Oxford Nanopore Ligation Sequencing Kit V14 (SQK-LSK114) and sequencing was done using a R10.4.1 PromethION flow cell.

Nanopore reads were assembled using Flye (2.9.1-b1780) (Kolmogorov et al., 2019) and https://github.com/fenderglass/Flye): “flye --nano-hq all.fastq.gz --scaffold --no-alt-contigs -- out-dir scaff_flye --threads 64” yielding a contig N50 of 205099. We aligned all larval RNA-seq reads (101361743 100bp paired end reads) to this assembly using STAR (v.2.7.10b) (Dobin et al., 2013). 86% of reads were uniquely mapped. A sorted BAM file of these alignments was used as input to Stringtie (v.2.1.0) (Pertea et al., 2015), with default parameters, to produce gene and transcript models. Protein sequences were predicted from these using Transdecoder (v.5.5.0), with the option to retain Pfam hits (http://transdecoder.github.io, (Haas et al., 2013)). For further protein level analyses such as phylogeny inference, we retained the longest transcript per gene, giving 29524 proteins. These data were 89% complete for BUSCO metazoa_odb9 (Simão et al., 2015).

### Orthology assignment and functional annotation

Orthologous pairs between all genes of our species of study were inferred from reciprocal best hits using the Smith-Waterman algorithm implemented in ssearch36 from the FASTA package (Pearson & Lipman, 1988).

For selected genes, orthology assignments were refined by phylogenetic analysis. Tree inference (as opposed to pairwise orthologs) for particular protein families used the method outlined in (Kapli et al., 2021). Alignments were constructed using programs from the HMMER3.3 package (http://hmmer.org/). Pfam hidden Markov models were searched against a protein database of representative metazoans (hmmsearch), using gathering threshold cutoffs (‘--cut_ga’), and sequences were aligned (hmmalign) and then filtered and trimmed (esl-alimask ‘--pavg 0.5’ and esl-alimanip ‘--minpp 0.3’, followed by ‘--lnfract 0.7’) on the basis of posterior probabilities of correct alignments and length of remaining sequence. Phylogenies were constructed using IQ-TREE, allowing the optimum model to be selected from the LG set (i.e., “-mset LG” in IQ-TREE).

For functional annotation, protein conserved domains were identified using hmmscan of the HMMER 3.3 package against the Pfam database (Finn et al., 2016). Analysis of GPCR and ion channels include genes annotated with the following Pfam domains: 7tm_1, 7tm_2, and 7tm_3 (for GPCR); and Ion_trans, Lig_chan, Neur_chan_memb, Neur_chan_LBD (for ion channels). Secreted proteins were annotated by presence of signal peptide using SignalP 5.0b (Almagro Armenteros et al., 2019).

### Fixation of specimens

For *In Situ* Hybridization (ISH), *Clytia* planulae were fixed in 3.7% formaldehyde/0.2% glutaraldehyde in PBS for 2 hours on ice, washed five times in PBST (PBS containing 0.1% Tween 20) for 10 minutes, then dehydrated in increasing concentrations of methanol in PBST (50%, 75%, 100%), and stored at -20°C. For Fluorescent ISH (FISH), Hybridization Chain Reaction (HCR) detection and immunostaining, *Clytia* planulae were fixed in a two step incubation with 1) HEM buffer (0.1 M HEPES pH 6.9, 50 mM EGTA, 10 mM MgSO4) containing 3.7% formaldehyde and 0.2% glutaraldehyde for 5 minutes at room temperature and 2) HEM buffer containing 3.7% formaldehyde for 2 hours at room temperature. After fixation, samples were washed 4 times for 5 minutes in PBST, dehydrated in increasing concentrations of methanol in PBST, and stored at -20°C.

*Astroides* and *Pocillopora* planulae were first relaxed using 0.4 M menthol in Millipore-Filtered (0.2 μm) Seawater (MFSW) on ice for 5 minutes. Then, larvae were incubated for 5 minutes at room temperature and fixed for 1 hour in formaldehyde 3.7% in MFSW buffered to pH 8 with HEPES buffer (on ice). After 4 rinses with PBST, specimens were dehydrated through a graded series of methanol in PBST and stored at -20°C in absolute methanol until use.

### Whole mount in situ hybridization (ISH), fluorescent in situ hybridization (FISH) and in situ hybridization chain reaction (HCR)

ISH was performed, as previously described (Lapébie et al., 2014) except that 4M Urea was used instead of 50% formamide in the hybridisation buffer (Sinigaglia et al., 2018). ISH probes were synthesized from either pGEM-T Easy plasmids (following insert amplification by PCR) or expressed sequence tag clones (Chevalier et al., 2006).

FISH was performed for xloc_042849 and pp2 using DIG-labeled probes. Signal development was carried out with an incubation overnight with a peroxidase-labeled anti-DIG antibody followed by washes in MABT (100 mM maleic acid pH 7.5, 150 mM NaCl, 0.1% Triton X-100). The fluorescence signal was developed using the TSA (Tyramide Signal Amplification) kit (TSA Plus Fluorescence Amplification kit, PerkinElmer, Waltham, MA) and Cy5 fluorophore (diluted 1/400 in TSA buffer: PBS/H2O20.0015%) at room temperature for 30 min. The development reaction was stopped by replacing the solution by PBST. After three washes for 5 minutes in PBST, nuclei were stained using 1:2000 Hoechst dye 33258 (1 mg/ml stock; Sigma-Aldrich) in PBST overnight at 4°C.

HCRs were performed for *Clytia* and *Astroides* planulae in eppendorf tubes using the Bruce et al., 2021 protocol (dx.doi.org/10.17504/protocols.io.bunznvf6). Probes for HCR were designed using the probe generator program created by Ryan Null (https://github.com/rwnull/insitu_probe_generator). Between 12 and 38 probe pairs using the full coding sequence as template were designed for each gene, ordered as oligo pools from IDT (Integrated DNA technologies) and suspended in TE buffer (10 mM Tris pH 8.0, 0.1 mM EDTA) at 1 μM. Fixed specimens kept at -20°C were progressively rehydrated in PBST and permeabilized in detergent solution (1.0% SDS, 0.5% Tween-20, 150 mM NaCl, 1 mM EDTA (pH 8), 50 mM Tris-HCl at pH 7.5) for 1 h. After four washes for 5 min in PBST at room temperature, the specimens were pre-hybridized in hybridization buffer (Molecular Instruments) for 1 h at 37 °C. The probes were then added to the hybridization buffer at a final concentration of 0.02 μM and the samples were let hybridize at 37 °C overnight with gentle agitation. After hybridization, the samples were washed four times for 15 min in probe wash buffer (Molecular instruments) at 37 °C, followed by two washes in 5× saline sodium citrate buffer containing 0.1% Tween-20 (5x SSCT) for 5 min, and two washes in 1x SSCT for 5 min at room temperature. They were then pre-amplified in amplification buffer (Molecular Instruments) for 30 min at room temperature. Meanwhile, H1 and H2 components of the HCR hairpins B1, B2 coupled to either Alexa546 or Alexa647 fluorophores (Molecular Instruments) were incubated separately at 95 °C for 90 s, cooled down to room temperature in the dark and then pooled together in the amplification buffer at a final concentration of 60 nM. The amplification was carried out overnight, in the dark and at room temperature. Excess hairpins were removed through two 5-minute washes and two 30-minute washes in 1× SSCT, followed by two 5-minute washes in PBST. Nuclei were stained using 1:2000 Hoechst dye 33258 (1 mg/ml stock; Sigma-Aldrich) in PBST overnight at 4°C.

### Immunostaining

Fixed specimens were gradually rehydrated from 100% methanol to PBST. *Clytia* larvae were extracted by performing two washes with PBS-Triton 0.2% for 10 minutes on rotation. *A. calycluaris* larvae were not extracted and instead were washed twice in PBST for 10 minutes on rotation. Samples were blocked in 0.2 μm-filtered blocking solution (3% Bovine Serum Albumin BSA in PBST) for one hour at room temperature (on rotation). After blocking, they were incubated overnight at 4°C in the primary antibody solution. The primary antibodies used in this study were: a mouse anti-LWamide antibody used at 1/20 (kindly provided by O. Koizumi (Koizumi et al., 2015)); a custom made, affinity purified rabbit polyclonal anti-QYFamide peptide antibody (produced by Covalab, Lyon) used at 13.5ng/ml in blocking solution; and a custom made affinity purified rat polyclonal anti-GLWamide antibody (Covalab) used at 2.45ng/ml in blocking solution. Following primary antibody incubation, samples were washed five times for 30 min each in PBST and incubated with fluorescence-conjugated secondary antibodies diluted 1:200 (anti-mouse Ig, anti-rat Ig, anti-rabbit Ig; Jackson Labs) and Hoechst dye 33342 1:2000 in PBST overnight at 4°C. Samples were then washed twice in PBST for 10 minutes and twice in 1x PBS for 5 minutes prior to mounting.

### Microscopy

Colorimetric ISH samples were mounted in 50% glycerol and imaged using a Zeiss Axio Imager 2 microscope. Images were processed with ImageJ v2.3.0/1.53f (Schindelin et al., 2012). For fluorescently labeled samples (FISH, HCR and immunohistochemistry), *Clytia* samples were mounted in Citifluor antifade mountant (Citifluor-EMS) and *Astroides* samples were dehydrated through a glycerol series and mounted in 90% glycerol/PBS 1x. Images were acquired using Leica Stellaris 5 or Sp8 laser scanning confocal microscopes and Z-stack maximum intensity projections were generated using ImageJ software.

### Settlement assays

To test the effect of different molecules on *Clytia* settlement, 4-well 1 mL plastic culture dishes (Thermo Fisher) were conditioned to generate a settlement-inducing biofilm. Culture dishes were left incubating in a tank of the IMEV/EMBRC-Fr Aquarium Service with a continuous flow of natural seawater containing different types of algae and microorganisms for a week. After this time period a biofilm was generated on the surfaces of the wells which efficiently induces *Clytia* planula settlement after overnight incubation (>90% of settlement). Treatments were performed in conditioned culture wells with each well containing 100 or 50 late 2-day planulae (52 hpf). Taurine, Glutamate, Betaine, GABA, Baclofen, Homotaurine (Sigma-Aldrich), were dissolved in non-filtered artificial seawater (ASW) at concentrations of 10-3, 10-4, 10-5 M. Each replicate contained a control condition with larvae incubated in non-filtered ASW. Settlement was scored about 16-20 hours after the start of the incubation. For *Astroides*, incubations were in 24-well plastic dishes containing 5 planulae per well. Settlement was scored after one week. Taurine and Glutamate were dissolved in natural seawater at concentrations of 10-3, 10-4, 10-5 M and 10-6M. Each replicate included a control condition with larvae incubated in natural seawater only. Solutions were replaced daily to compensate for molecule degradation. To assess the rate of settlement, settled larvae were counted daily.

## Acknowledgements

We thank Marie-Jeanne Arguel at the France Genomix platform (IPMC Sophia Antipolis) for assistance with single-cell sequencing and *Astroides* genome sequencing; Sophie Brasseur, Marc Liebeck and Tom Tarpey for their participation in larval settlement assays; Alice Mirasole and Antonia Chiarore for assistance in the field, and Chloé Carbonne for the photo in Figure 1A and her assistance with the aquarium facilities. We thank the Oceanopolis Brest tropical aquarium scientific director Pierre Ternat for providing access to *Pocillopora acuta* colonies and their planulae, and Aurélien Morellon and Maureen Midol for help with monitoring of planulation events and care of the parent colonies. We thank Thomas Lamonerie, the Houliston/Momose and Copley group members for helpful discussions. *Clytia* were provided by the Marine Resources Centre and microscopy facilities by the Imaging Platform (PIM) of Institut de la Mer de Villefranche, both supported by EMBRC-France. The French state funds of EMBRC-France are managed by the ANR within the investments of the Future program. This research was supported by the Sorbonne-Université programme Emergence grant 2021-2023 ‘CLASSE’, the French ANR grant 2022-2026 ‘MOPSEA’, the H2020/Marie Skłodowska-Curie ITN “EvoCell” Grant agreement no. 766053, and the French Government through the National Research Agency—Investments for the Future (‘4Oceans-Make Our Planet Great Again’ grant, ANR-17-MOPGA-0001).

## Author contributions

E.H. and R.R.C. conceived the study. J.R.-M., E.H. and R.R.C. designed the experiments and wrote the manuscript. J.R.-M. and A.F. performed the single-cell experiments. J.R.-M. performed the bulk RNA-seq experiments, genome extraction for *Astroides*, in situ hybridizations, immunostainings, and settlement assays. J.R.-M. and R.R.C. performed bioinformatics analysis. N.T. and I.D.-C. provided the *Astroides* and *Pocillopora* planulae. J.R.-M., E.H. and R.R.C. analyzed and interpreted the data. E.H., R.R.C, I.D.-C. and N.T. acquired the funding. All authors contributed to manuscript revisions.

## References

1. Albrecht, J., & Schousboe, A. (2005). Taurine interaction with neurotransmitter receptors in the CNS: an update. Neurochemical Research, 30(12), 1615–1621.

2. Almagro Armenteros, J. J., Tsirigos, K. D., Sønderby, C. K., Petersen, T. N., Winther, O., Brunak, S., von Heijne, G., & Nielsen, H. (2019). SignalP 5.0 improves signal peptide predictions using deep neural networks. Nature Biotechnology, 37(4), 420–423.

3. Anderson, P. A. V., & Trapido-Rosenthal, H. G. (2009). Physiological and chemical analysis of neurotransmitter candidates at a fast excitatory synapse in the jellyfish Cyanea capillata (Cnidaria, Scyphozoa). Invertebrate Neuroscience: IN, 9(3-4), 167–173.

4. Arendt, D., Tosches, M. A., & Marlow, H. (2016). From nerve net to nerve ring, nerve cord and brain--evolution of the nervous system. Nature Reviews. Neuroscience, 17(1), 61–72.

5. Attenborough, R. M. F., Hayward, D. C., Wiedemann, U., Forêt, S., Miller, D. J., & Ball, E. E. (2019). Expression of the neuropeptides RFamide and LWamide during development of the coral Acropora millepora in relation to settlement and metamorphosis. Developmental Biology, 446(1), 56–67.

6. Berking, S. (1986). Is homarine a morphogen in the marine hydroid Hydractinia? Roux’s Archives of Developmental Biology: The Official Organ of the EDBO, 195(1), 33–38.

7. Berking, S. (1988). Taurine found to stabilize the larval state is released upon induction of metamorphosis in the hydrozon Hydractinia. Roux’s Archives of Developmental Biology: The Official Organ of the EDBO, 197(6), 321–327.

8. Biggers, W. J., Pires, A., Pechenik, J. A., Johns, E., Patel, P., Polson, T., & Polson, J. (2012). Inhibitors of nitric oxide synthase induce larval settlement and metamorphosis of the polychaete annelid Capitella teleta. Invertebrate Reproduction & Development, 56(1), 1–13.

9. Birch, S., & Plachetzki, D. (2023). Multisensory integration by polymodal sensory neurons dictates larval settlement in a brainless cnidarian larva. Molecular Ecology, 32(14), 3892– 3907.

10. Bishop, C. D., & Brandhorst, B. P. (2001). NO/cGMP signaling and HSP90 activity represses metamorphosis in the sea urchin Lytechinus pictus. The Biological Bulletin, 201(3), 394– 404.

11. Bishop, C. D., Pires, A., Norby, S.-W., Boudko, D., Moroz, L. L., & Hadfield, M. G. (2008). Analysis of nitric oxide-cyclic guanosine monophosphate signaling during metamorphosis of the nudibranch Phestilla sibogae Bergh (Gastropoda: Opisthobranchia). Evolution & Development, 10(3), 288–299.

12. Bodo, F., & Bouillon, J. (1968). Étude histologique du développement embryonnaire de quelques hydroméduses de Roscoff: Phialidium hemisphaericum (L.), Obelia sp. Péron et Lesueur, Sarsia eximia (Allman), Podocoryne carnea (Sars), Gonionemus vertens Agassiz. Cahiers de Biologie Marine.

13. Bosch, T. C. G. (2009). Hydra and the evolution of stem cells. *BioEssays: News and Reviews in Molecular*, Cellular and Developmental Biology, 31(4), 478–486.

14. Brown, B. E., & Bythell, J. C. (2005). Perspectives on mucus secretion in reef corals. Marine Ecology Progress Series, 296, 291–309.

15. Carbonne, C., Comeau, S., Chan, P. T. W., Plichon, K., Gattuso, J.-P., & Teixidó, N. (2022). Early life stages of a Mediterranean coral are vulnerable to ocean warming and acidification. Biogeosciences, 19(19), 4767–4777.

16. Castañeda, O., Sotolongo, V., Amor, A. M., Stöcklin, R., Anderson, A. J., Harvey, A. L., Engström, A., Wernstedt, C., & Karlsson, E. (1995). Characterization of a potassium channel toxin from the Caribbean Sea anemone Stichodactyla helianthus. Toxicon: Official Journal of the International Society on Toxinology, 33(5), 603–613.

17. Cavalier-Smith, T. (2017). Origin of animal multicellularity: precursors, causes, consequences— the choanoflagellate/sponge transition, neurogenesis and the Cambrian explosion. Philosophical Transactions of the Royal Society of London. Series B, Biological Sciences, 372(1713), 20150476.

18. Chari, T., Weissbourd, B., Gehring, J., Ferraioli, A., Leclère, L., Herl, M., Gao, F., Chevalier, S., Copley, R. R., Houliston, E., Anderson, D. J., & Pachter, L. (2021). Whole-animal multiplexed single-cell RNA-seq reveals transcriptional shifts across Clytia medusa cell types. Science Advances, 7(48), eabh1683.

19. Chevalier, S., Martin, A., Leclère, L., Amiel, A., & Houliston, E. (2006). Polarised expression of FoxB and FoxQ2 genes during development of the hydrozoan Clytia hemisphaerica. Development Genes and Evolution, 216(11), 709–720.

20. Chiori, R., Jager, M., Denker, E., Wincker, P., Da Silva, C., Le Guyader, H., Manuel, M., & Quéinnec, E. (2009). Are Hox genes ancestrally involved in axial patterning? Evidence from the hydrozoan Clytia hemisphaerica (Cnidaria). PloS One, 4(1), e4231.

21. Columbus-Shenkar, Y. Y., Sachkova, M. Y., Macrander, J., Fridrich, A., Modepalli, V., Reitzel, A. M., Sunagar, K., & Moran, Y. (2018). Dynamics of venom composition across a complex life cycle. eLife, 7.

22. Comes, S., Locascio, A., Silvestre, F., d’Ischia, M., Russo, G. L., Tosti, E., Branno, M., & Palumbo, A. (2007). Regulatory roles of nitric oxide during larval development and metamorphosis in Ciona intestinalis. Developmental Biology, 306(2), 772–784.

23. Croce, J., Duloquin, L., Lhomond, G., McClay, D. R., & Gache, C. (2006). Frizzled5/8 is required in secondary mesenchyme cells to initiate archenteron invagination during sea urchin development. Development, 133(3), 547–557.

24. Cunning, R., Bay, R. A., Gillette, P., Baker, A. C., & Traylor-Knowles, N. (2018). Comparative analysis of the Pocillopora damicornis genome highlights role of immune system in coral evolution. Scientific Reports, 8(1), 16134.

25. Darras, S., Gerhart, J., Terasaki, M., Kirschner, M., & Lowe, C. J. (2011). β-catenin specifies the endomesoderm and defines the posterior organizer of the hemichordate Saccoglossus kowalevskii. Development, 138(5), 959–970.

26. Dobin, A., Davis, C. A., Schlesinger, F., Drenkow, J., Zaleski, C., Jha, S., Batut, P., Chaisson, M., & Gingeras, T. R. (2013). STAR: ultrafast universal RNA-seq aligner. Bioinformatics, 29(1), 15–21.

27. Ferraioli, A. (2022). Comparison of cell types across life cycle stages of the hydrozoan Clytia hemisphaerica. PhD thesis, *Sorbonne université*.

28. Finn, R. D., Coggill, P., Eberhardt, R. Y., Eddy, S. R., Mistry, J., Mitchell, A. L., Potter, S. C., Punta, M., Qureshi, M., Sangrador-Vegas, A., Salazar, G. A., Tate, J., & Bateman, A. (2016). The Pfam protein families database: towards a more sustainable future. Nucleic Acids Research, 44(D1), D279–D285.

29. García-Castro, H., Kenny, N. J., Iglesias, M., Álvarez-Campos, P., Mason, V., Elek, A., Schönauer, A., Sleight, V. A., Neiro, J., Aboobaker, A., Permanyer, J., Irimia, M., Sebé-Pedrós, A., & Solana, J. (2021). ACME dissociation: a versatile cell fixation-dissociation method for single-cell transcriptomics. Genome Biology, 22(1), 89.

30. Gerhart, J., Lowe, C., & Kirschner, M. (2005). Hemichordates and the origin of chordates. Current Opinion in Genetics & Development, 15(4), 461–467.

31. Gilbert, E., Craggs, J., & Modepalli, V. (2024). Gene Regulatory Network that Shaped the Evolution of Larval Apical Organ in Cnidaria. Molecular Biology and Evolution, 41(1).

32. Gilbert, E., Teeling, C., Lebedeva, T., Pedersen, S., Chrismas, N., Genikhovich, G., & Modepalli, V. (2022). Molecular and cellular architecture of the larval sensory organ in the cnidarian Nematostella vectensis. Development, 149(16).

33. Gilis, M., Meibom, A., Domart-Coulon, I., Grauby, O., Stolarski, J., & Baronnet, A. (2014). Biomineralization in newly settled recruits of the scleractinian coral Pocillopora damicornis. Journal of Morphology, 275(12), 1349–1365.

34. Gröger, H., & Schmid, V. (2001). Larval development in Cnidaria: a connection to Bilateria? Genesis, 29(3), 110–114.

35. Haas, B. J., Papanicolaou, A., Yassour, M., Grabherr, M., Blood, P. D., Bowden, J., Couger, M. B., Eccles, D., Li, B., Lieber, M., MacManes, M. D., Ott, M., Orvis, J., Pochet, N., Strozzi, F., Weeks, N., Westerman, R., William, T., Dewey, C. N., … Regev, A. (2013). De novo transcript sequence reconstruction from RNA-seq using the Trinity platform for reference generation and analysis. Nature Protocols, 8(8), 1494–1512.

36. Hayakawa, E., Watanabe, H., Menschaert, G., Holstein, T. W., Baggerman, G., & Schoofs, L. (2019). A combined strategy of neuropeptide prediction and tandem mass spectrometry identifies evolutionarily conserved ancient neuropeptides in the sea anemone Nematostella vectensis. PloS One, 14(9), e0215185.

37. Houliston, E., Momose, T., & Manuel, M. (2010). Clytia hemisphaerica: a jellyfish cousin joins the laboratory. Trends in Genetics: TIG, 26(4), 159–167.

38. Hunnekuhl, V. S., & Akam, M. (2014). An anterior medial cell population with an apical-organ-like transcriptional profile that pioneers the central nervous system in the centipede Strigamia maritima. Developmental Biology, 396(1), 136–149.

39. Huxtable, R. J. (1992). Physiological actions of taurine. Physiological Reviews, 72(1), 101–163.

40. Islam, M. S., Leissing, T. M., Chowdhury, R., Hopkinson, R. J., & Schofield, C. J. (2018). 2-Oxoglutarate-dependent oxygenases. Annual Review of Biochemistry, 87, 585–620.

41. Kaminow, B., Yunusov, D., & Dobin, A. (2021). STARsolo: accurate, fast and versatile mapping/quantification of single-cell and single-nucleus RNA-seq data. In bioRxiv (p. 2021.05.05.442755).

42. Kapli, P., Flouri, T., & Telford, M. J. (2021). Systematic errors in phylogenetic trees. Current Biology: CB, 31(2), R59–R64.

43. Kayal, E., Bentlage, B., Sabrina Pankey, M., Ohdera, A. H., Medina, M., Plachetzki, D. C., Collins, A. G., & Ryan, J. F. (2018). Phylogenomics provides a robust topology of the major cnidarian lineages and insights on the origins of key organismal traits. BMC Evolutionary Biology, 18(1).

44. Khadka, A., Martínez-Bartolomé, M., Burr, S. D., & Range, R. C. (2018). A novel gene’s role in an ancient mechanism: secreted Frizzled-related protein 1 is a critical component in the anterior-posterior Wnt signaling network that governs the establishment of the anterior neuroectoderm in sea urchin embryos. EvoDevo, 9, 1.

45. Koizumi, O., Hamada, S., Minobe, S., Hamaguchi-Hamada, K., Kurumata-Shigeto, M., Nakamura, M., & Namikawa, H. (2015). The nerve ring in cnidarians: its presence and structure in hydrozoan medusae. Zoology, 118(2), 79–88.

46. Kolmogorov, M., Yuan, J., Lin, Y., & Pevzner, P. A. (2019). Assembly of long, error-prone reads using repeat graphs. Nature Biotechnology, 37(5), 540–546.

47. Kopp, C., Domart-Coulon, I., Barthelemy, D., & Meibom, A. (2016). Nutritional input from dinoflagellate symbionts in reef-building corals is minimal during planula larval life stage. Science Advances, 2(3), e1500681.

48. Korsunsky, I., Millard, N., Fan, J., Slowikowski, K., Zhang, F., Wei, K., Baglaenko, Y., Brenner, M., Loh, P.-R., & Raychaudhuri, S. (2019). Fast, sensitive and accurate integration of single-cell data with Harmony. Nature Methods, 16(12), 1289–1296.

49. Krasovec, G., Pottin, K., Rosello, M., Quéinnec, É., & Chambon, J.-P. (2021). Apoptosis and cell proliferation during metamorphosis of the planula larva of Clytia hemisphaerica (Hydrozoa, Cnidaria). Developmental Dynamics: An Official Publication of the American Association of Anatomists, 250(12), 1739–1758.

50. Lambert, I. H., Kristensen, D. M., Holm, J. B., & Mortensen, O. H. (2015). Physiological role of taurine--from organism to organelle. *Acta Physiologica (Oxford*, England), 213(1), 191–212.

51. Lapébie, P., Ruggiero, A., Barreau, C., Chevalier, S., Chang, P., Dru, P., Houliston, E., & Momose, T. (2014). Differential responses to Wnt and PCP disruption predict expression and developmental function of conserved and novel genes in a cnidarian. PLoS Genetics, 10(9), e1004590.

52. Layden, M. J., Boekhout, M., & Martindale, M. Q. (2012). Nematostella vectensis achaete-scute homolog NvashA regulates embryonic ectodermal neurogenesis and represents an ancient component of the metazoan neural specification pathway. Development, 139(5), 1013– 1022.

53. Lechable, M., Jan, A., Duchene, A., Uveira, J., Weissbourd, B., Gissat, L., Collet, S., Gilletta, L., Chevalier, S., Leclere, L., Peron, S., Barreau, C., Lasbleiz, R., Houliston, E., & Momose, T. (2020). An improved whole life cycle culture protocol for the hydrozoan genetic model Clytia hemisphaerica. Biology Open, 9(11), bio057034.

54. Leclère, L., Horin, C., Chevalier, S., Lapébie, P., Dru, P., Peron, S., Jager, M., Condamine, T., Pottin, K., Romano, S., Steger, J., Sinigaglia, C., Barreau, C., Quiroga Artigas, G., Ruggiero, A., Fourrage, C., Kraus, J. E. M., Poulain, J., Aury, J.-M., … Copley, R. R. (2019). The genome of the jellyfish Clytia hemisphaerica and the evolution of the cnidarian life-cycle. Nature Ecology & Evolution, 3(5), 801–810.

55. Leitz, T., & Lay, M. (1995). Metamorphosin A is a neuropeptide. Roux’s Archives of Developmental Biology: The Official Organ of the EDBO, 204(4), 276–279.

56. Lemaître, Q. I. B., Bartsch, N., Kouzel, I. U., Busengdal, H., Richards, G. S., Steinmetz, P. R. H., & Rentzsch, F. (2023). NvPrdm14d-expressing neural progenitor cells contribute to non-ectodermal neurogenesis in Nematostella vectensis. Nature Communications, 14(1), 4854.

57. Levy, S., Brekhman, V., Bakhman, A., Malik, A., Sebé-Pedrós, A., Kosloff, M., & Lotan, T. (2021). Ectopic activation of GABAB receptors inhibits neurogenesis and metamorphosis in the cnidarian Nematostella vectensis. Nature Ecology & Evolution, 5(1), 111–121.

58. Levy, S., Elek, A., Grau-Bové, X., Menéndez-Bravo, S., Iglesias, M., Tanay, A., Mass, T., & Sebé-Pedrós, A. (2021). A stony coral cell atlas illuminates the molecular and cellular basis of coral symbiosis, calcification, and immunity. Cell, 184(11), 2973–2987.e18.

59. Love, M. I., Huber, W., & Anders, S. (2014). Moderated estimation of fold change and dispersion for RNA-seq data with DESeq2. Genome Biology, 15(12), 550.

60. Lowe, C. J., Wu, M., Salic, A., Evans, L., Lander, E., Stange-Thomann, N., Gruber, C. E., Gerhart, J., & Kirschner, M. (2003). Anteroposterior patterning in hemichordates and the origins of the chordate nervous system. Cell, 113(7), 853–865.

61. Marlow, H., Tosches, M. A., Tomer, R., Steinmetz, P. R., Lauri, A., Larsson, T., & Arendt, D. (2014). Larval body patterning and apical organs are conserved in animal evolution. BMC Biology, 12, 7.

62. Massé, A. (2018). Association between the microboring chlorophyte of the genus Ostreobium and scleractinian corals. PhD thesis, *Sorbonne université*.

63. Monier, M., Nuez, I., Borne, F., & Courtier-Orgogozo, V. (2024). Higher evolutionary dynamics of gene copy number for Drosophila glue genes located near short repeat sequences. BMC Ecology and Evolution, 24(1), 18.

64. Moran, Y., Praher, D., Schlesinger, A., Ayalon, A., Tal, Y., & Technau, U. (2013). Analysis of soluble protein contents from the nematocysts of a model sea anemone sheds light on venom evolution. Marine Biotechnology, 15(3), 329–339.

65. Müller, W. A., & Leitz, T. (2002). Metamorphosis in the Cnidaria. Canadian Journal of Zoology, 80(10), 1755–1771.

66. Müller, W. A., Mitze, A., Wickhorst, J.-P., & Meier-Menge, H. M. (1977). Polar morphogenesis in early hydroid development: Action of caesium, of neurotransmitters and of an intrinsic head activator on pattern formation. Wilhelm Roux’s Archives of Developmental Biology, 182(4), 311–328.

67. Nakanishi, N., Yuan, D., Jacobs, D. K., & Hartenstein, V. (2008). Early development, pattern, and reorganization of the planula nervous system in Aurelia (Cnidaria, Scyphozoa). Development Genes and Evolution, 218(10), 511–524.

68. Pani, A. M., Mullarkey, E. E., Aronowicz, J., Assimacopoulos, S., Grove, E. A., & Lowe, C. J. (2012). Ancient deuterostome origins of vertebrate brain signalling centres. Nature, 483(7389), 289–294.

69. Pasantes-Morales, H. (1996). Volume regulation in brain cells: cellular and molecular mechanisms. Metabolic Brain Disease, 11(3), 187–204.

70. Pearson, W. R., & Lipman, D. J. (1988). Improved tools for biological sequence comparison. Proceedings of the National Academy of Sciences of the United States of America, 85(8), 2444–2448.

71. Pertea, M., Pertea, G. M., Antonescu, C. M., Chang, T.-C., Mendell, J. T., & Salzberg, S. L. (2015). StringTie enables improved reconstruction of a transcriptome from RNA-seq reads. Nature Biotechnology, 33(3), 290–295.

72. Piraino, S., Zega, G., Di Benedetto, C., Leone, A., Dell’Anna, A., Pennati, R., Carnevali, D. C., Schmid, V., & Reichert, H. (2011). Complex neural architecture in the diploblastic larva of Clava multicornis (Hydrozoa, Cnidaria). The Journal of Comparative Neurology, 519(10), 1931–1951.

73. Poustka, A. J., Kühn, A., Groth, D., Weise, V., Yaguchi, S., Burke, R. D., Herwig, R., Lehrach, H., & Panopoulou, G. (2007). A global view of gene expression in lithium and zinc treated sea urchin embryos: new components of gene regulatory networks. Genome Biology, 8(5), R85.

74. Rentzsch, F., Fritzenwanker, J. H., Scholz, C. B., & Technau, U. (2008). FGF signalling controls formation of the apical sensory organ in the cnidarian Nematostella vectensis. Development, 135(10), 1761–1769.

75. Ryan, J. F., Mazza, M. E., Pang, K., Matus, D. Q., Baxevanis, A. D., Martindale, M. Q., & Finnerty, J. R. (2007). Pre-bilaterian origins of the Hox cluster and the Hox code: evidence from the sea anemone, Nematostella vectensis. PloS One, 2(1), e153.

76. Sabin, K. Z., Chen, S., Hill, E. M., Weaver, K. J., Yonke, J., Kirkman, M., Redwine, W. B., Klompen, A. M. L., Zhao, X., Guo, F., McKinney, M. C., Dewey, J. L., & Gibson, M. C. (2024). Graded FGF activity patterns distinct cell types within the apical sensory organ of the sea anemone Nematostella vectensis. Developmental Biology, 510, 50–65.

77. Sachkova, M. Y., Landau, M., Surm, J. M., Macrander, J., Singer, S. A., Reitzel, A. M., & Moran, Y. (2020). Toxin-like neuropeptides in the sea anemone Nematostella unravel recruitment from the nervous system to venom. Proceedings of the National Academy of Sciences of the United States of America, 117(44), 27481–27492.

78. Santagata, S., Resh, C., Hejnol, A., Martindale, M. Q., & Passamaneck, Y. J. (2012). Development of the larval anterior neurogenic domains of Terebratalia transversa (Brachiopoda) provides insights into the diversification of larval apical organs and the spiralian nervous system. EvoDevo, 3, 3.

79. Saransaari, P., & Oja, S. S. (2000). Taurine and neural cell damage. Amino Acids, 19(3-4), 509– 526.

80. Schindelin, J., Arganda-Carreras, I., Frise, E., Kaynig, V., Longair, M., Pietzsch, T., Preibisch, S., Rueden, C., Saalfeld, S., Schmid, B., Tinevez, J.-Y., White, D. J., Hartenstein, V., Eliceiri, K., Tomancak, P., & Cardona, A. (2012). Fiji: an open-source platform for biological-image analysis. Nature Methods, 9(7), 676–682.

81. Schmich, J., Trepel, S., & Leitz, T. (1998). The role of GLWamides in metamorphosis of Hydractinia echinata. Development Genes and Evolution, 208(5), 267–273.

82. Schultz, D. T., Haddock, S. H. D., Bredeson, J. V., Green, R. E., Simakov, O., & Rokhsar, D. S. (2023). Ancient gene linkages support ctenophores as sister to other animals. Nature, 618(7963), 110–117.

83. Sebé-Pedrós, A., Saudemont, B., Chomsky, E., Plessier, F., Mailhé, M.-P., Renno, J., Loe-Mie, Y., Lifshitz, A., Mukamel, Z., Schmutz, S., Novault, S., Steinmetz, P. R. H., Spitz, F., Tanay, A., & Marlow, H. (2018). Cnidarian Cell Type Diversity and Regulation Revealed by Whole-Organism Single-Cell RNA-Seq. Cell, 173(6), 1520–1534.e20.

84. Siebert, S., Farrell, J. A., Cazet, J. F., Abeykoon, Y., Primack, A. S., Schnitzler, C. E., & Juliano, C. E. (2019). Stem cell differentiation trajectories in Hydra resolved at single-cell resolution. Science, 365(6451).

85. Simão, F. A., Waterhouse, R. M., Ioannidis, P., Kriventseva, E. V., & Zdobnov, E. M. (2015). BUSCO: assessing genome assembly and annotation completeness with single-copy orthologs. Bioinformatics, 31(19), 3210–3212.

86. Sinigaglia, C., Busengdal, H., Leclère, L., Technau, U., & Rentzsch, F. (2013). The bilaterian head patterning gene six3/6 controls aboral domain development in a cnidarian. PLoS Biology, 11(2), e1001488.

87. Sinigaglia, C., Busengdal, H., Lerner, A., Oliveri, P., & Rentzsch, F. (2015). Molecular characterization of the apical organ of the anthozoan Nematostella vectensis. Developmental Biology, 398(1), 120–133.

88. Sinigaglia, C., Thiel, D., Hejnol, A., Houliston, E., & Leclère, L. (2018). A safer, urea-based in situ hybridization method improves detection of gene expression in diverse animal species. Developmental Biology, 434(1), 15–23.

89. Song, H., Hewitt, O. H., & Degnan, S. M. (2021). Arginine Biosynthesis by a Bacterial Symbiont Enables Nitric Oxide Production and Facilitates Larval Settlement in the Marine-Sponge Host. Current Biology: CB, 31(2), 433–437.e3.

90. Steger, J., Cole, A. G., Denner, A., Lebedeva, T., Genikhovich, G., Ries, A., Reischl, R., Taudes, E., Lassnig, M., & Technau, U. (2022). Single-cell transcriptomics identifies conserved regulators of neuroglandular lineages. Cell Reports, 40(12), 111370.

91. Steinmetz, P. R., Urbach, R., Posnien, N., Eriksson, J., Kostyuchenko, R. P., Brena, C., Guy, K., Akam, M., Bucher, G., & Arendt, D. (2010). Six3 demarcates the anterior-most developing brain region in bilaterian animals. EvoDevo, 1(1), 14.

92. Steinworth, B. M., Martindale, M. Q., & Ryan, J. F. (2023). Gene Loss may have Shaped the Cnidarian and Bilaterian Hox and ParaHox Complement. Genome Biology and Evolution, 15(1).

93. Takahashi, T., & Hatta, M. (2011). The Importance of GLWamide Neuropeptides in Cnidarian Development and Physiology. Journal of Amino Acids, 2011, 424501.

94. Takahashi, T., & Takeda, N. (2015). Insight into the molecular and functional diversity of cnidarian neuropeptides. International Journal of Molecular Sciences, 16(2), 2610–2625.

95. Takeda, N., Kon, Y., Quiroga Artigas, G., Lapébie, P., Barreau, C., Koizumi, O., Kishimoto, T., Tachibana, K., Houliston, E., & Deguchi, R. (2018). Identification of jellyfish neuropeptides that act directly as oocyte maturation-inducing hormones. Development, 145(2).

96. Thomas, M. B., Freeman, G., & Martin, V. J. (1987). The embryonic origin of neurosecretory cells and the role of nerve cells in metamorphosis in Clytia.

97. Traag, V. A., Waltman, L., & van Eck, N. J. (2019). From Louvain to Leiden: guaranteeing well-connected communities. Scientific Reports, 9(1), 5233.

98. Ueda, N., Richards, G. S., Degnan, B. M., Kranz, A., Adamska, M., Croll, R. P., & Degnan, S. M. (2016). An ancient role for nitric oxide in regulating the animal pelagobenthic life cycle: evidence from a marine sponge. Scientific Reports, 6, 37546.

99. Vandermeulen, J. (1974). Studies on reef corals. II. Fine structure of planktonic planula larva of Pocillopora damicornis, with emphasis on the aboral epidermis. Marine Biology, 27, 239– 249.

100. Walther, M., Ulrich, R., Kroiher, M., & Berking, S. (1996). Metamorphosis and pattern formation in Hydractinia echinata, a colonial hydroid. The International Journal of Developmental Biology, 40(1), 313–322.

101. Wolf, F. A., Angerer, P., & Theis, F. J. (2018). SCANPY: large-scale single-cell gene expression data analysis. Genome Biology, 19(1), 15.

102. Yankura, K. A., Martik, M. L., Jennings, C. K., & Hinman, V. F. (2010). Uncoupling of complex regulatory patterning during evolution of larval development in echinoderms. BMC Biology, 8, 143.

103. Zhang, Y., He, L.-S., Zhang, G., Xu, Y., Lee, O.-O., Matsumura, K., & Qian, P.-Y. (2012). The regulatory role of the NO/cGMP signal transduction cascade during larval attachment and metamorphosis of the barnacle Balanus (=Amphibalanus) amphitrite. The Journal of Experimental Biology, 215(Pt 21), 3813–3822.

