## Supplementary Figures for "Aboral cell types of *Clytia* and coral larvae have shared features and link taurine to the regulation of settlement"

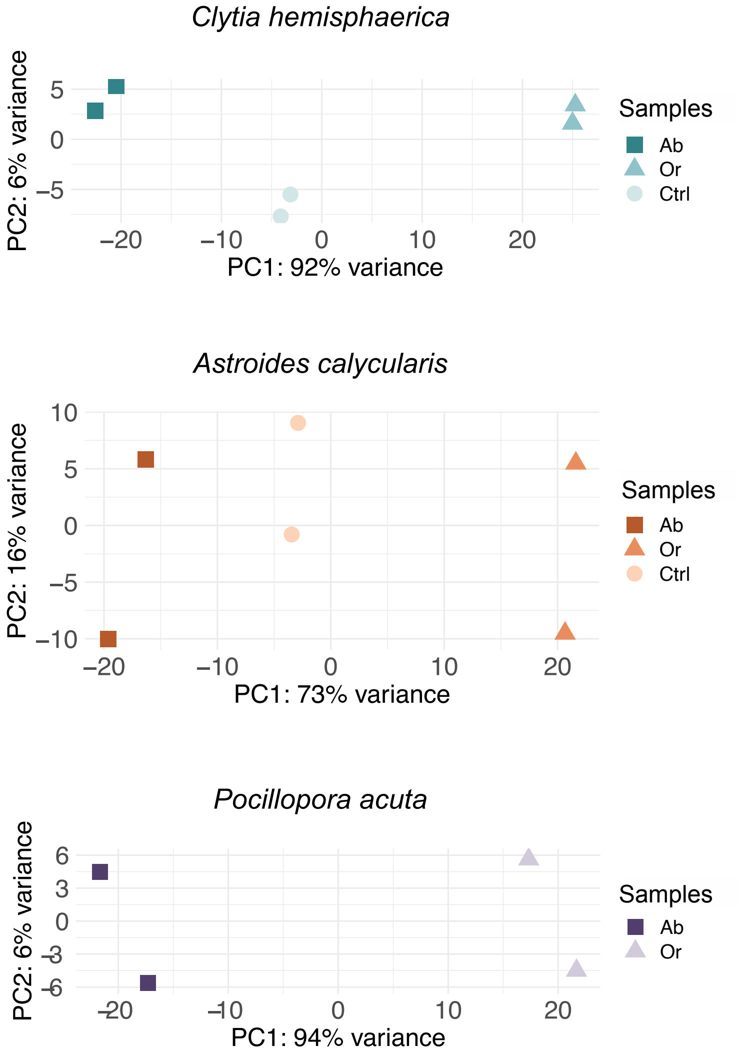


**Supplementary Figure 1: Principal Component Analysis (PCA) of planula Aboral/Oral transcriptomes.**

PCA plots of RNA-seq libraries for each species generated using normalized counts (see Methods). Samples for *Clytia* (top) and *Astroides* (middle) are aboral (Ab), oral (Or), and uncut control (Ctrl); for *Pocillopora* (bottom), the samples are aboral (Ab) and oral (Or).

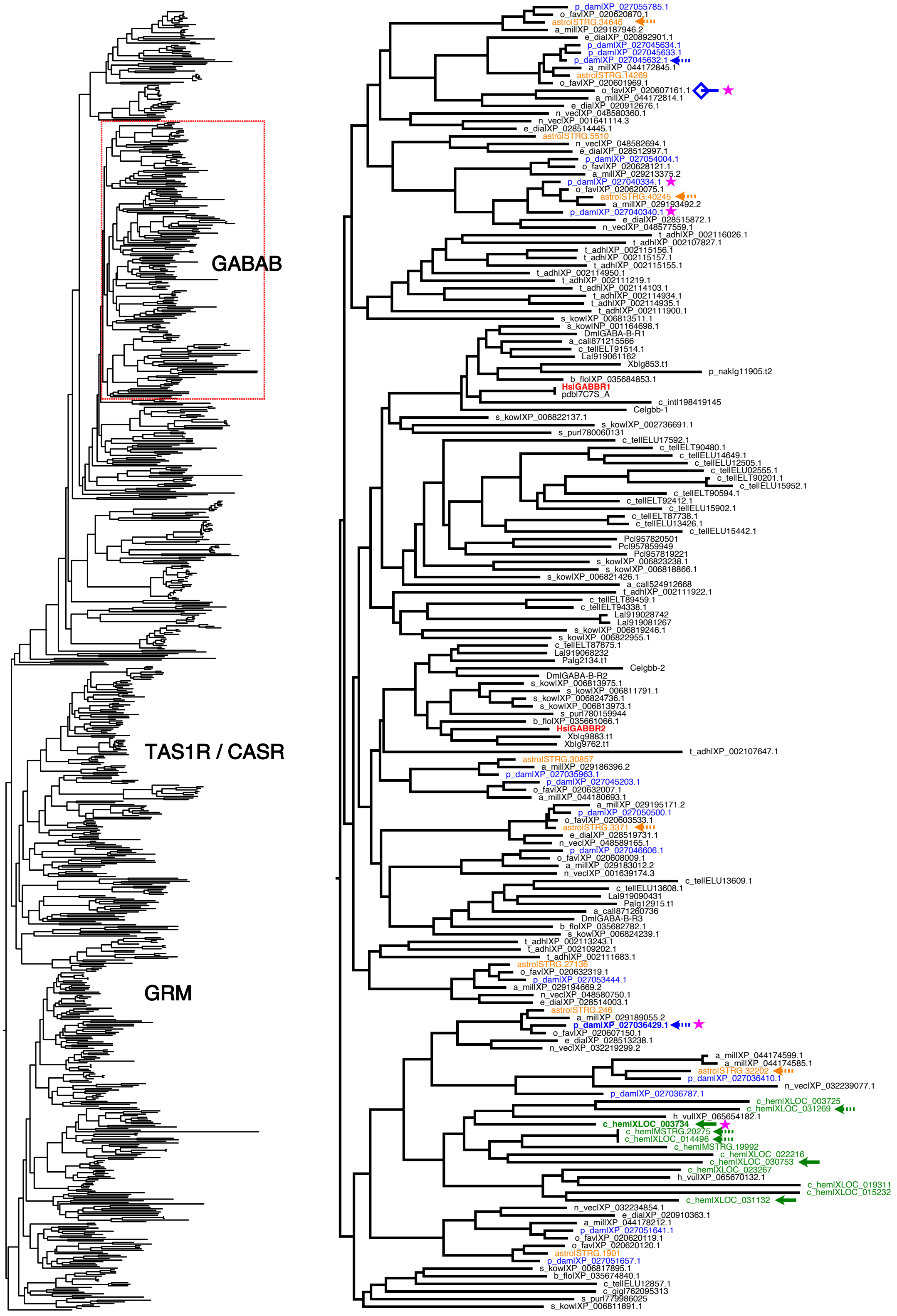
GABAB
TAS1R / CASR
GRM
HsGABBR1
HsGABBR2

**Supplementary Figure 2: Maximum Likelihood phylogenetic analysis of metabotropic-type GPCRs.**

The tree on the left includes putative metabotropic receptor sequences categorized by including the Pfam domains ANF_receptor and 7tm_3. GABAB: Metabotropic GABA receptor; TAS1R/CASR: Taste Receptors type 1/Calcium-Sensing Receptors; GRM: Metabotropic Glutamate receptors. The tree on the right is a zoomed view of the region outlined by the dotted red box. Sequences from *Clytia*, *Astroides*, and *Pocillopora* are highlighted in green, orange, and blue, respectively, and human sequences are marked in red. Arrows indicate aboral-enriched sequences (continuous lines lfc>1, dotted lines lfc>0). Magenta stars indicate expression in GLWamide cells. The blue square indicates the expected placement of the *Pocillopora* aboral-enriched gene XP_027050808.1, which is missing the ANF_receptor domain in its sequence, based on reciprocal best hit analysis. p_dam = *Pocillopora damicornis*; o_fav = *Orbicella faveolata*; astro = *Astroides calycularis*; a_mil = *Acropora millepora*; e_dia = *Exaiptasia diaphana*; n_vec = *Nematostella vectensis*; t_adh = *Trichoplax adhaerens*; s_kow = *Saccoglossus kowalevskii*; Dm = *Drosophila melanogaster*; a_cal = *Aplysia californica*; c_tel = *Capitella teleta*; La = *Lingula anatina*; Xb = *Xenoturbella bockii*; p_nak = *Praesagittifera naikaiensis*; b_flo = *Branchiostoma floridae*; Hs = *Homo sapiens*; pdb = Protein Data Bank (3D structures); c_int = *Ciona intestinalis*; Ce = *Caenorhabditis elegans*; s_pur = *Strongylocentrotus purpuratus*; Pc = *Priapulus caudatus*; Pa = *Phoronis australis*; c_hem = *Clytia hemisphaerica*; h_vul = *Hydra vulgaris*.

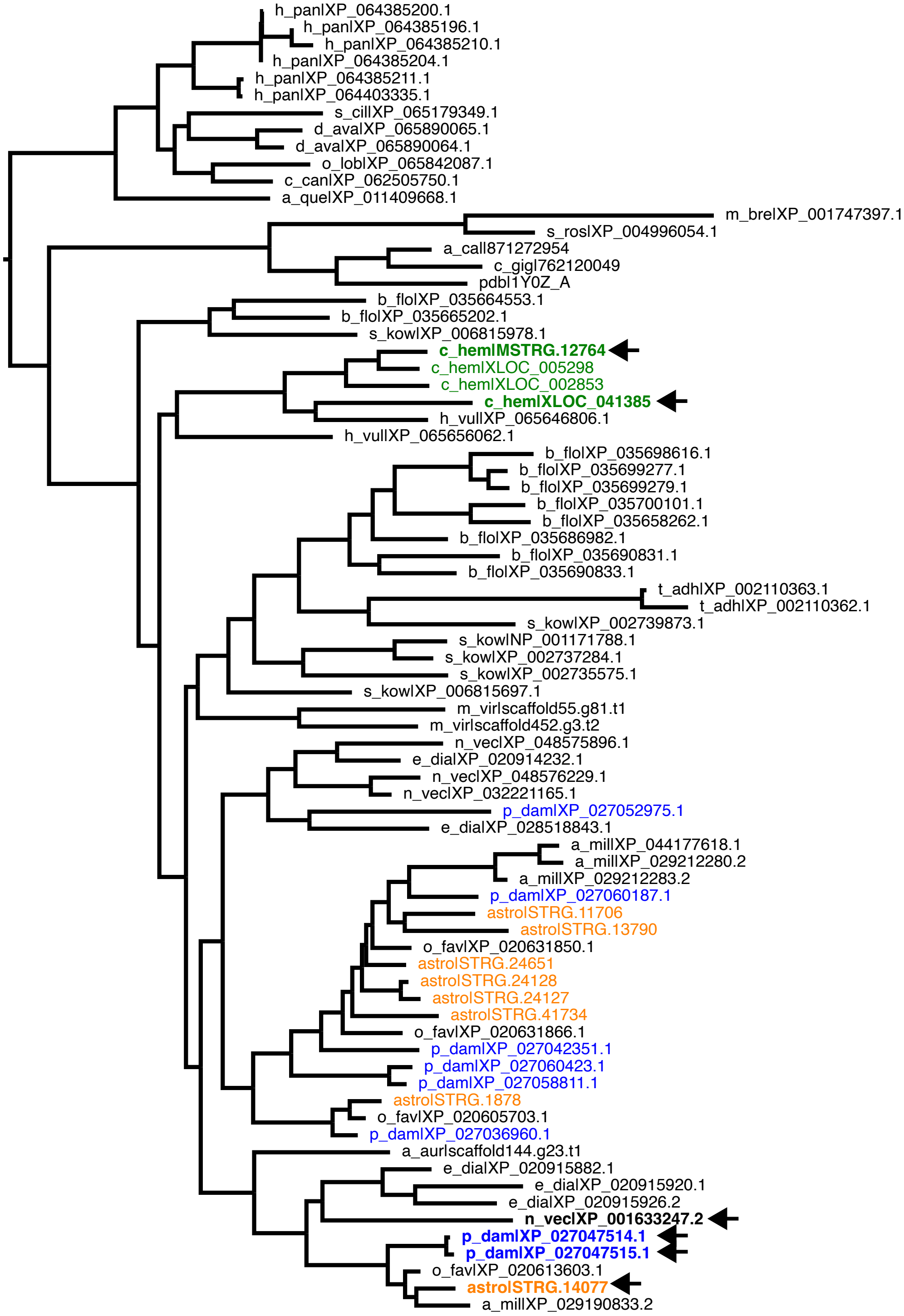

h_pan|XP_064385200.1
h_pan|XP_064385196.1
h_pan|XP_064385210.1
h_pan|XP_064385204.1
h_pan|XP_064385211.1
h_pan|XP_064403335.1
s_cil|XP_065179349.1
d_ava|XP_065890065.1
d_ava|XP_065890064.1
o_lob|XP_065842087.1
c_can|XP_062505750.1
a_que|XP_011409668.1
m_bre|XP_001747397.1
s_ros|XP_004996054.1
a_cal|871272954
c_gig|762120049
pdb|1Y0Z_A
b_flo|XP_035664553.1
b_flo|XP_035665202.1
s_kow|XP_006815978.1
c_hem|MSTRG.12764
c_hem|XLOC_005298
c_hem|XLOC_002853
c_hem|XLOC_041385
h_vul|XP_065646806.1
h_vul|XP_065656062.1
b_flo|XP_035698616.1
b_flo|XP_035699277.1
b_flo|XP_035699279.1
b_flo|XP_035700101.1
b_flo|XP_035658262.1
b_flo|XP_035686982.1
b_flo|XP_035690831.1
b_flo|XP_035690833.1
t_adh|XP_002110363.1
t_adh|XP_002110362.1
s_kow|XP_002739873.1
s_kow|NP_001171788.1
s_kow|XP_002737284.1
s_kow|XP_002735575.1
s_kow|XP_006815697.1
m_vir|scaffold55.g81.t1
m_vir|scaffold452.g3.t2
n_vec|XP_048575896.1
e_dia|XP_020914232.1
n_vec|XP_048576229.1
n_vec|XP_032221165.1
p_dam|XP_027052975.1
e_dia|XP_028518843.1
a_mil|XP_044177618.1
a_mil|XP_029212280.2
a_mil|XP_029212283.2
p_dam|XP_027060187.1
astro|STRG.11706
astro|STRG.13790
o_fav|XP_020631850.1
astro|STRG.24651
astro|STRG.24128
astro|STRG.24127
astro|STRG.41734
o_fav|XP_020631866.1
p_dam|XP_027042351.1
p_dam|XP_027060423.1
p_dam|XP_027058811.1
astro|STRG.1878
o_fav|XP_020605703.1
p_dam|XP_027036960.1
a_aur|scaffold144.g23.t1
e_dia|XP_020915882.1
e_dia|XP_020915920.1
e_dia|XP_020915926.2
n_vec|XP_001633247.2
p_dam|XP_027047514.1
p_dam|XP_027047515.1
o_fav|XP_020613603.1
astro|STRG.14077
a_mil|XP_029190833.2

**Supplementary Figure 3: Maximum Likelihood phylogenetic analysis of TauD proteins.** Sequences from *Clytia*, *Astroides*, and *Pocillopora* are highlighted in green, orange, and blue, respectively. Black arrows indicate expression enriched in the aboral end. The *Nematostella* sequence is equivalent to the one in (Sinigaglia et al., 2015). h_pan = *Halichondria panicea*; s_cil = *Sycon ciliatum*; d_ava = *Dysiea avara;* o_lob = *Oscarella lobularis*; c_can =*Corticium candelabrum*; a_que = *Amphimedon queenslandica*; m_bre = *Monosiga brevicollis*; s_ros = *Salpingoeca rosetta*; a_cal = *Aplysia californica*; c_gig = Crassostrea gigas; pdb = Protein Data Bank (3D structures); b_flo = *Branchiostoma floridae*; s_kow = *Saccoglossus kowalevskii*; c_hem = *Clytia hemisphaerica*; h_vul = *Hydra vulgaris*; t_adh = *Trichoplax adhaerens*; m_vir = *Morbakka virulenta*; N_vec = *Nematostella vectensis;* e_dial = *Exaiptasia diaphana*; p_dam = *Pocillopora damicornis*; a_mil = *Acropora millepora*; astro = *Astroides calycularis*; o_fav = *Orbicella faveolata*; a_aur = *Aurelia aurita*.

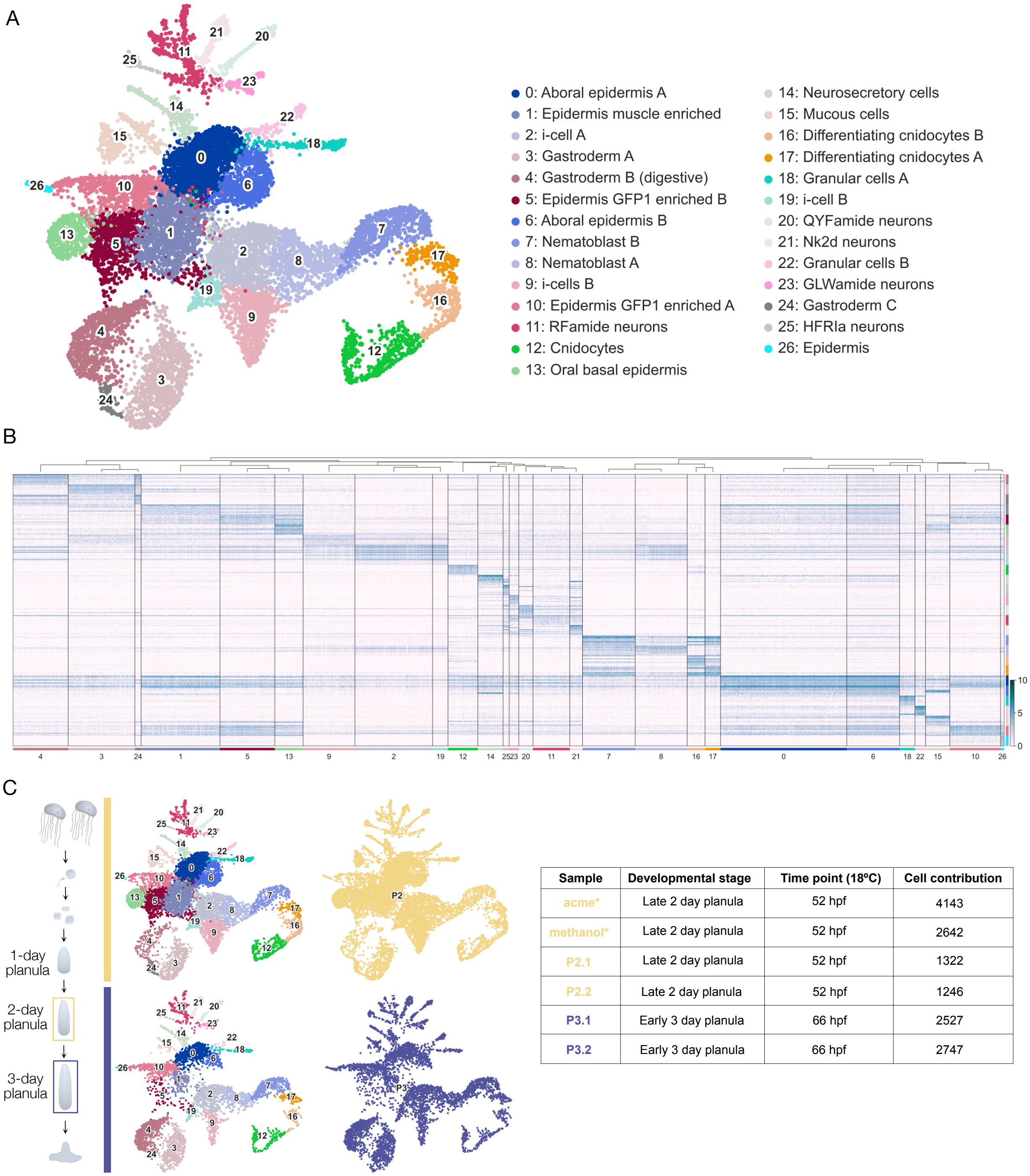


| Sample | Developmental stage | Time point (18ºC) | Cell contribution |
|---|---|---|---|
| acme* | Late 2 day planula | 52 hpf | 4143 |
| methanol* | Late 2 day planula | 52 hpf | 2642 |
| P2.1 | Late 2 day planula | 52 hpf | 1322 |
| P2.2 | Late 2 day planula | 52 hpf | 1246 |
| P3.1 | Early 3 day planula | 66 hpf | 2527 |
| P3.2 | Early 3 day planula | 66 hpf | 2747 |

**Supplementary Figure 4: Cluster annotation and sample contribution for the *Clytia* planula scRNAseq dataset.**

(A) UMAP representation of *Clytia* planula scRNAseq data with all 27 clusters labeled by cell type identity. (B) Heatmap of the 10 top marker genes for each cluster. Cluster color labels correspond to (A). (C) On the left, UMAP representation of cells sampled from 2-day (top) and

3-day planulae (bottom). Left plots are labeled by cell cluster and right plots are labeled by developmental stage (color code matching to the schematics on the left). On the right, a table with information for each library contributing to the dataset.

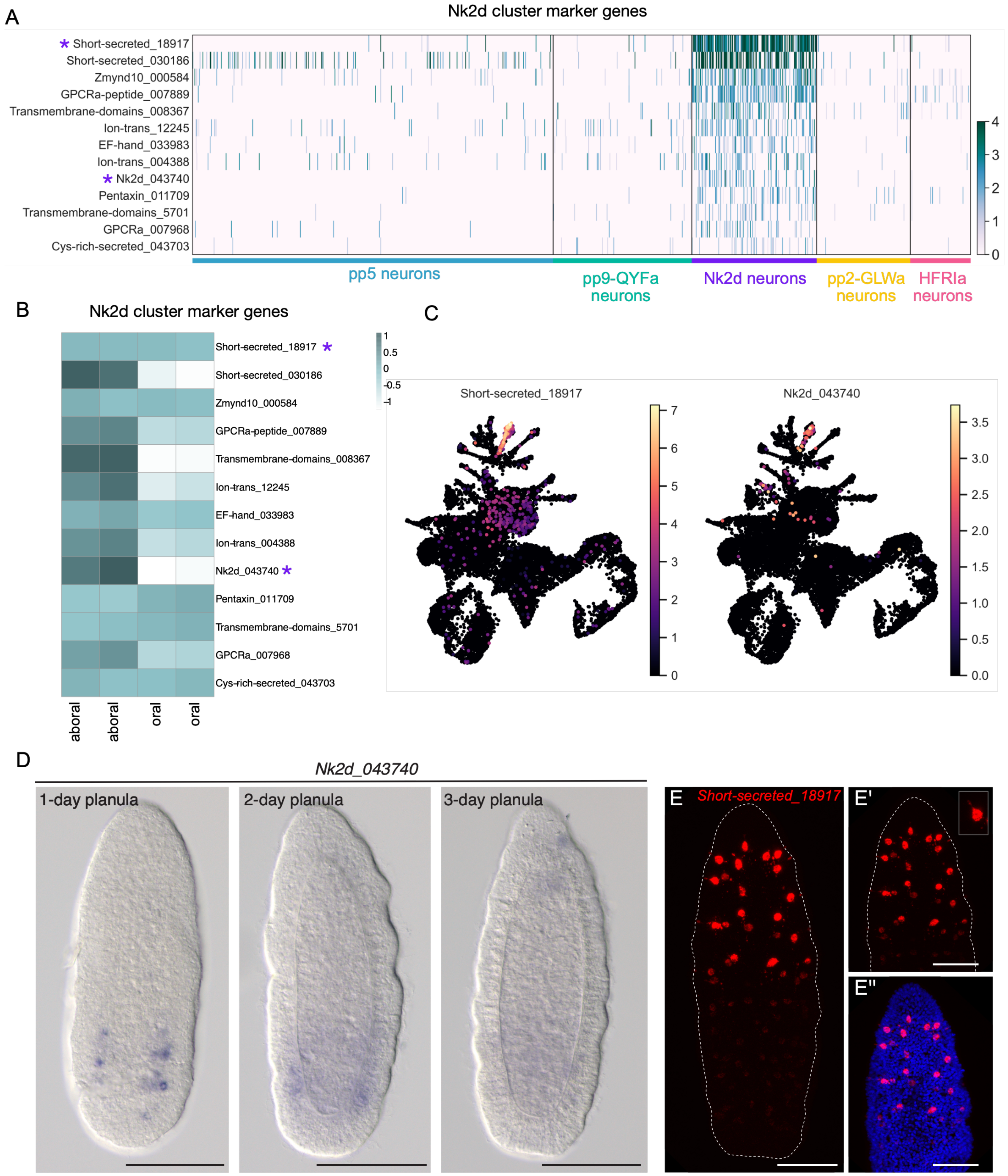

**Supplementary Figure 5: Characterization of the “Nk2d neural” cluster.**

(A) Heatmap of the marker genes for the Nk2d cell cluster. Purple asterisks indicate the genes shown in (C, D, E-E’’). Only neural clusters are included. Cell cluster annotation labels correspond to Fig. 2C and Supplementary Fig. 4. (B) Heatmap of aboral/oral expression (normalized counts) for the genes shown in (A). The “Nk2d cell” cluster marker genes included a mixture of aboral-enriched genes and genes with no aboral or oral enrichment. (C) UMAP plots of an aboral enriched marker gene (Nk2d_043740) and a homogeneously expressed marker gene (Short-secreted_18917). Expression of these two genes maps to subpopulations within the cluster. (D) ISH for Nk2d_043740 in 1-day, 2-day planula and 3-day planula stages. Expression is detected at the 1-day stage in cells at the base of the epidermis at the aboral end. Fainter signal was seen in the same area at 2-day planula. (E) HCR for Short-secreted_18917 in a 2-day planula showing expression in ganglionic-like neurons brightly stained in the oral half. Panels on the right (E’, E’’) show a magnification of the oral end. Inset shows close-up of an individual cell body. Short-secreted_18917 (red) and Hoechst dye (blue). Planulae all oriented with the oral pole uppermost. Scale bars, 100 µm.

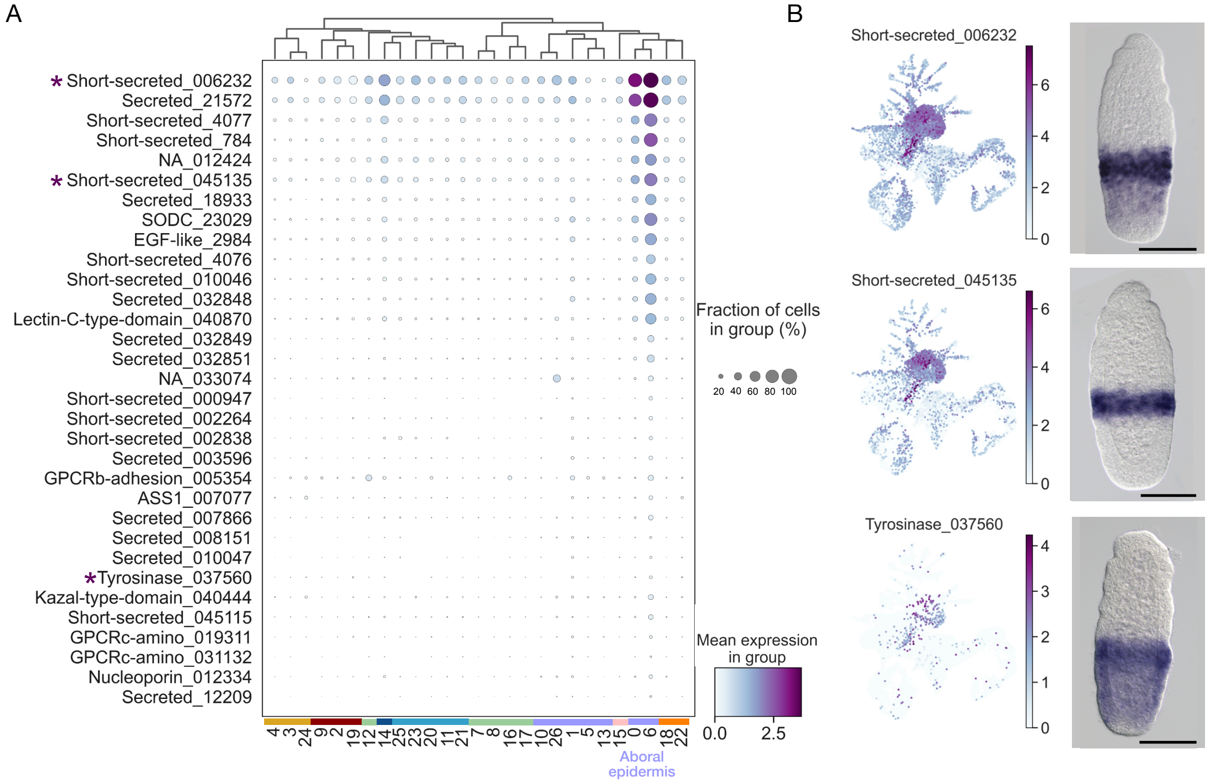


**Supplementary Figure 6: Putative aboral epidermis clusters in the *Clytia* planula cell atlas.**

(A) Gene expression dot plot of specific markers for the epidermal clusters 0 and 6. Gene IDs are listed on the left and cluster identities are indicated at the bottom. The color coding for the clusters corresponds to the cell class annotations in Fig. 2. The color gradient indicates average expression, and the size of the dots indicates the percentage of cells expressing the gene. Purple asterisks indicate the genes shown in (B). Note that the cluster markers include several short secreted proteins, a putative neuropeptide (Che-pp6 which is predicted to generate SRLLFamide peptides) and two class C GPCRs. (B) UMAP plots for three markers and corresponding in situ hybridization patterns in 2-day planula stage. Scale bars, 100 µm.

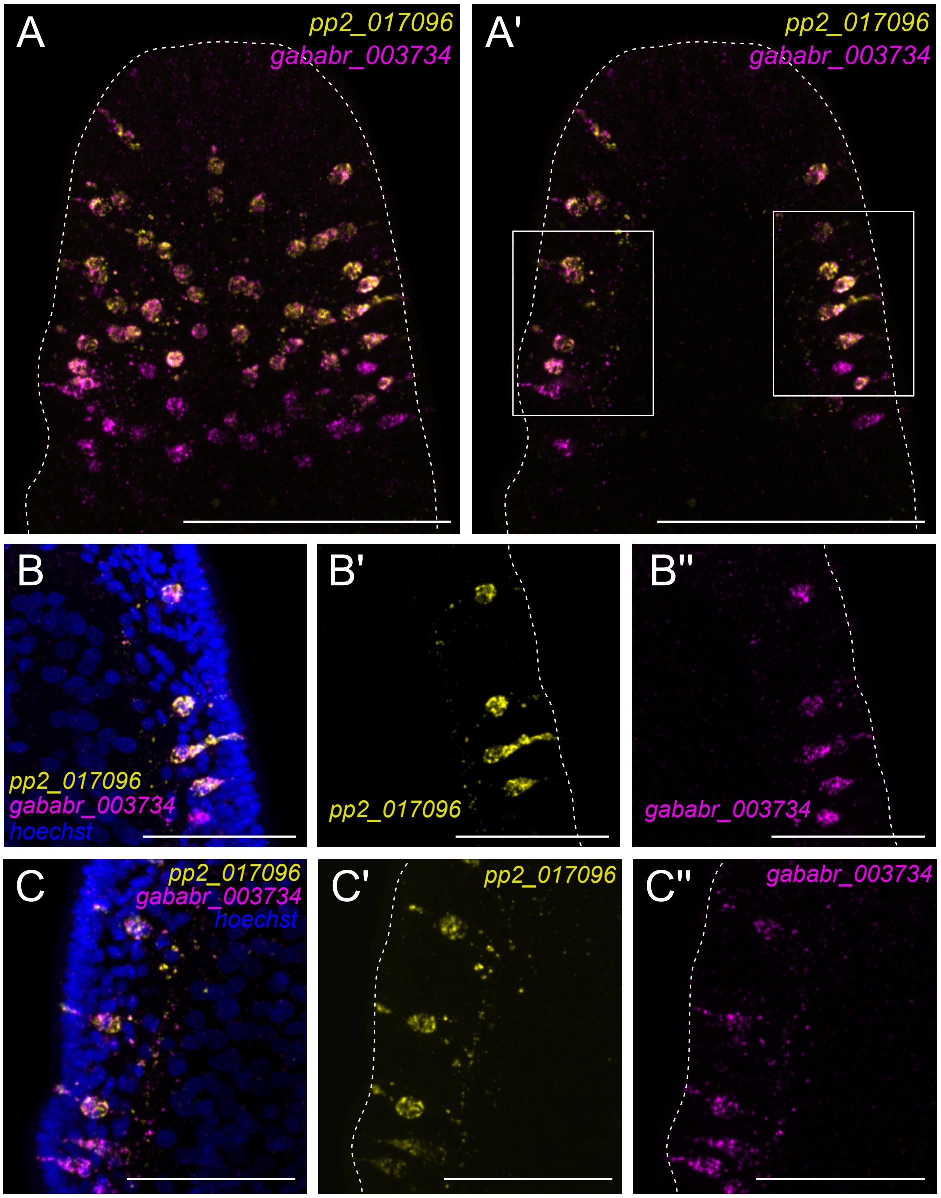


**Supplementary Figure 7: Coexpression of the GLWamide precursor pp2 and GABA$_B$R-like in aboral cells of the *Clytia* planula.**

Double FISH showing co-expression of pp2 (yellow) and GABA$_B$R (magenta) mRNAs in the same cells of the aboral epidermis. (A and A') Maximum intensity projections of confocal Z-stacks (A) and selected stacks from the medial region (A') of the aboral pole. Scale bars, 100 µm. (B-C'') Maximum intensity projections of confocal Z-stacks of outlined zones in (A'): (B-B'') corresponds to the square on the right and (C-C'') to the square on the left. pp2 (yellow), GABA$_B$R (magenta) and Hoechst dye (blue). Single channels are shown in (B', B'' and C', C''). Scale bars, 50 µm.

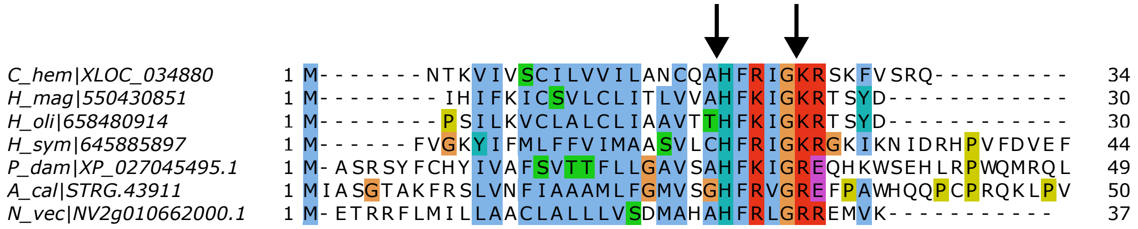


**Supplementary Figure 8: Putative HFRIamide neuropeptide in Cnidaria.**

Alignment of the first 50 amino acids of HFRIamide protein precursors from selected of hydrozoan and anthozoan species. The arrows mark the likely signal peptide cleavage site (left) and neuropeptide cleavage site (right). C_hem = *Clytia hemisphaerica*; H_mag = *Hydra magnipapillata*; H_oli = *Hydra oligactis*; H_sym = *Hydractinia symbiolongicapus*; P_dam = *Pocillopora damicornis*; A_cal = *Astroides calycularis*; N_vec = *Nematostella vectensis*. Coloring conventions are those of Clustal (Larkin et al., 2007).

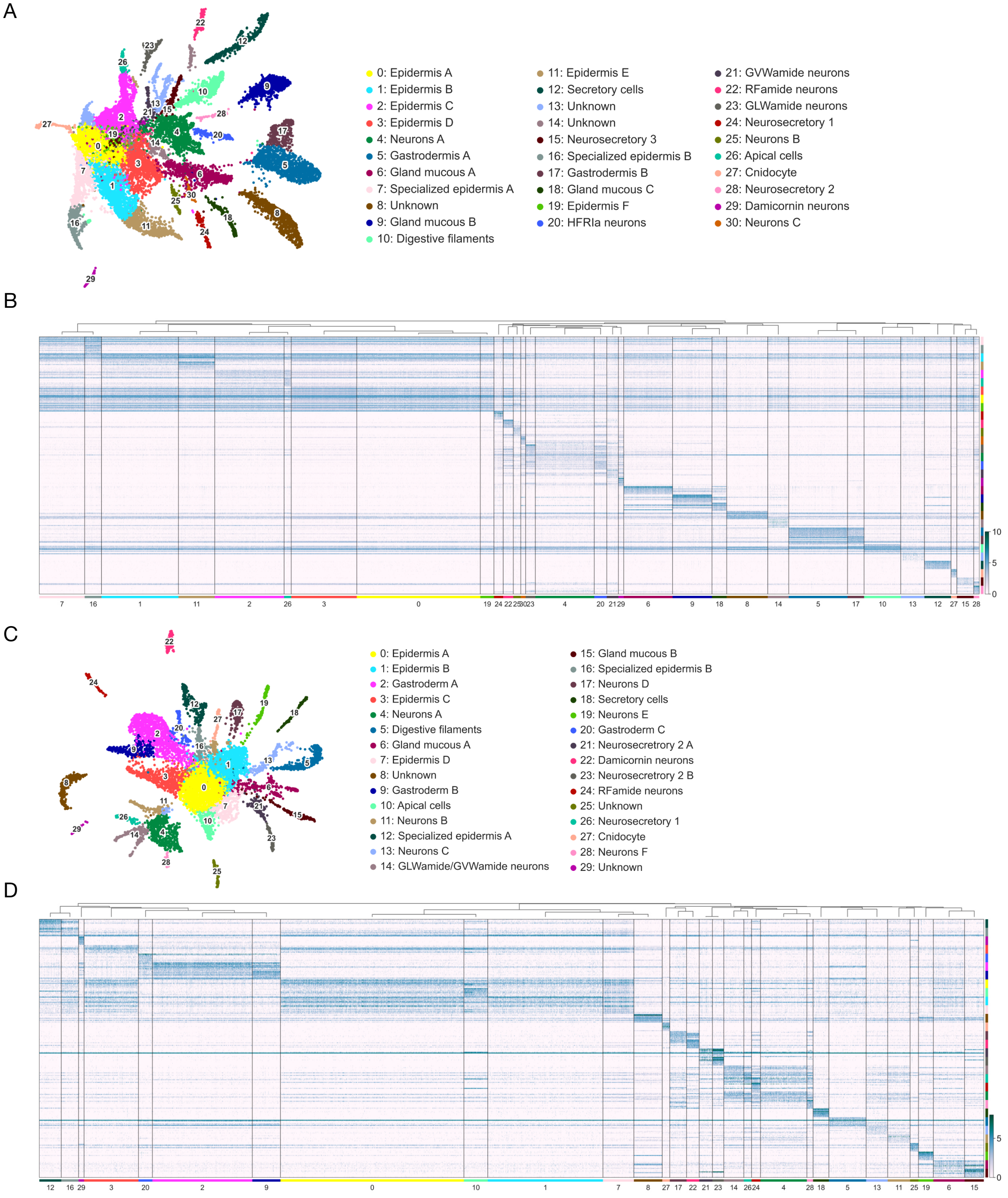


**Supplementary Figure 9: The *Astroides* and *Pocillopora* planula cell atlases.**

(A and C) UMAP representations of scRNAseq data from *Astroides* (A) and *Pocillopora* (C) planulae, with all clusters labeled by cell type identity. (B and D) Heatmaps of the 10 top marker genes for each cluster in *Astroides* and *Pocillopora*, respectively. The cluster color labels correspond to their respective atlases.

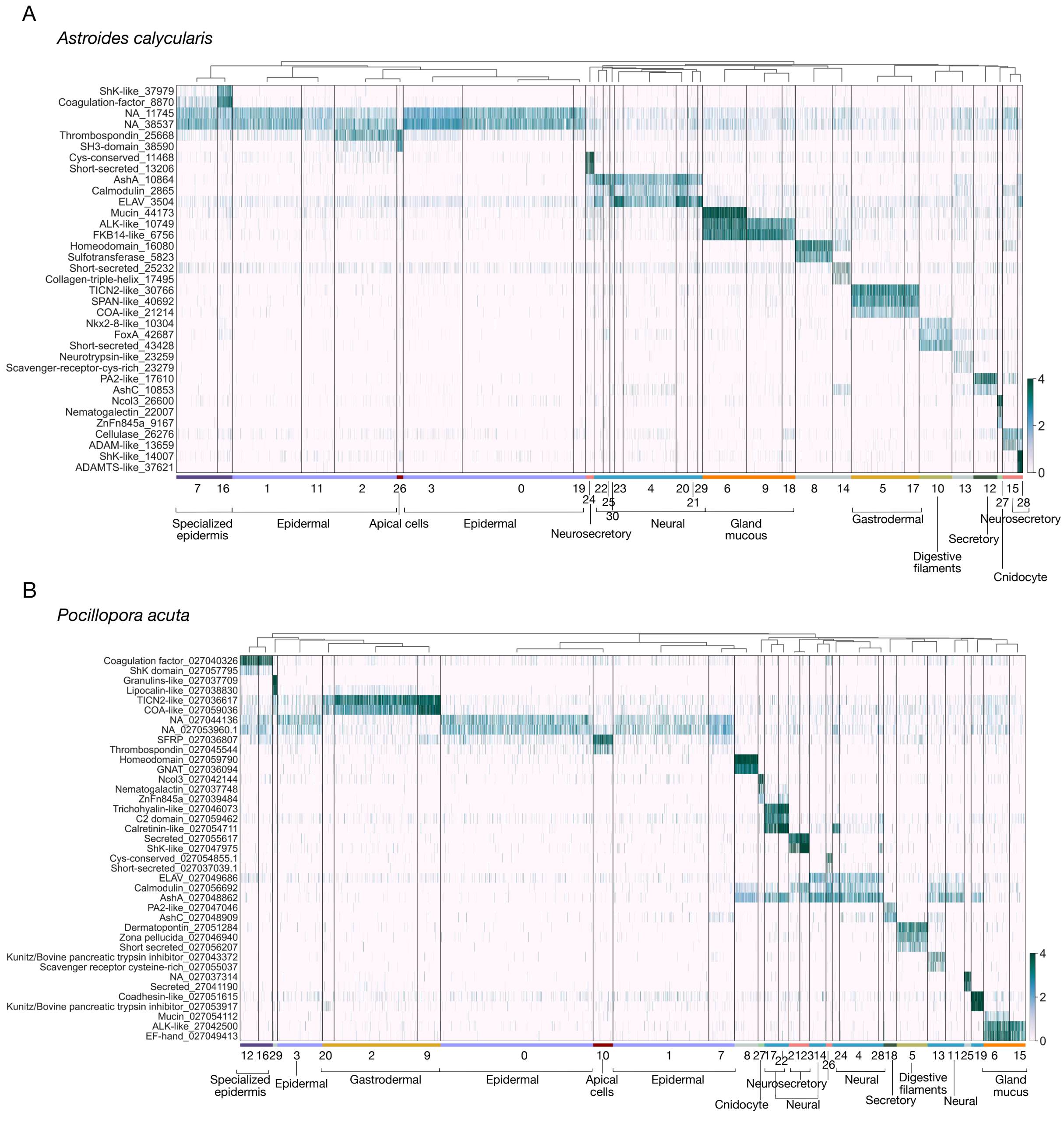


**Supplementary Figure 10: Annotation of coral planula cell atlases.**

Heatmap of diagnostic marker genes used to annotate cell type classes for *Astroides* (top) and *Pocillopora* (bottom) planula scRNAseq clusters. Gene names are taken from orthology assignments. In cases where assignment was not clear, the conserved protein domain is indicated. NA indicates absence of orthology assignment and conserved domains. Trailing numbers are unique gene identifiers from this project.

| GENERAL IDENTITY | CLUSTER *A. calycularis* | CLUSTER *P. acuta* | CELL TYPE | DIAGNOSTIC MARKERS | REFERENCE |
|---|---|---|---|---|---|
| Epidermal | 0, 1, 2, 3, 11, 19 | 0, 1, 3, 7 | | Sox3: STRG.11680 / XP_027038477.1 | *N. vectensis:* Steger et al., 2022 |
| | | | | STRG.11745 / XP_027044136.1 | |
| | | | | STRG.38537 / XP_027053960.1 | |
| Spezialized epidermis | 7, 16 | 12, 16 | | Shk domain-like: STRG.37979 / XP_027057795.1 | |
| | | | | STRG.8870 / XP_027040326.1 | |
| Apical cells | 26 | 10 | | ISX-like: STRG.15235 / XP_027046639.1 | *N. vectensis*: Gilbert et al., 2022 & Sabin et al., 2024 |
| | | | | FoxQ2a: STRG.21723 / XP_027053483.1 | *N. vectensis*: Sinigaglia et al., 2013 |
| | | | | FGF: STRG.38785 / XP_027049367.1 | *N. vectensis*: Rentzsch et al., 2008 |
| | | | | PoxA: STRG.27319 / XP_027045474.1 | *N. vectensis*: Gilbert et al., 2022 |
| Gastroderm | 5, 17 | 2, 9, 20 | | ZF-C3H11: STRG.10901 / XP_027055412.1 | *N. vectensis:* Steger et al., 2022 |
| | | | | Testican2-like: STRG.30766 / XP_027036617.1 | *N. vectensis:* Steger et al., 2022 |
| | | | | COA-like2: STRG.21214 / XP_027059036.1 | *N. vectensis:* Steger et al., 2022 |
| Digestive filaments | 10 | 5 | | Nkx2-8-like: STRG.10304 / XP_027038915.1 | *S. pistillata*: Levy et al., 2021 |
| | | | | OIT3-like12: STRG.21702 / XP_027046940.1 | *S. pistillata*: Levy et al., 2021 |
| Neural | 4, 20, 21, 22, 23, 25, 30, 29 | 4, 11, 13,14, 17, 19, 22, 24, 28 | | AshA: STRG.10864 / XP_027048862.1 | *N. vectensis*: Layden et al;, 2012 & Steger et al., 2022 |
| | | | | ELAV - STRG.3504 / XP_027049686.1 | |
| | 21 | 14 | GVWamide neurons | GVWamide: STRG.21557 / XP_027059473.1 | |
| | 22 | 24 | RFamide neurons | RFamide: STRG.11240 / XP_027051563.1 | |
| | 23 | 14 | GLWamide neurons | GLWamide: STRG.15971 / XP_027060553.1 | |
| | 29 | 22 | Damicornin neurons | Damicornin: STRG.4391 / XP_027051004.1 | |
| | 20 | - | HFRIa neurons | HFRIa: STRG.43911 / XP_027045495.1 | |
| Gland mucous | 6, 9, 18 | 6, 15 | | RFX4: STRG.29170 / XP_027047616.1 | *N. vectensis:* Steger et al., 2022 & Gilbert et al., 20222 |
| | | | | Mucin : STRG.44173 / XP_027054112.1 | *N. vectensis:* Steger et al., 2022 |
| | | | | ALK-like: STRG.10749 / XP_027042500.1 | *N. vectensis:* Steger et al., 2022 |
| Neurosecretory | 15, 24, 28 | 21, 23, 26 | | | |
| | 24 | 26 | Neurosecretory 1 | Secreted, Cys-conserved: STRG.11468 / XP_027054855.1 | |
| | | | | Nkx2.5: STRG.11493 / XP_027041575.1 | |
| | 28 | 21, 23 | Neurosecretory 2 | Shk domain-like: STRG.14007 / XP_027047975.1 | |
| | | | | Pax6: STRG.28117 / XP_027036201.1 | |
| Secretory | 12 | 18 | | AshC: STRG.10853 / XP_027048909.1 | *N. vectensis:* Steger et al., 2022 |
| | | | | PA2-like2: STRG.17610 / XP_027047046.1 | |
| Cnidocyte | 27 | 27 | | Znfn845: STRG.9167 / XP_027039484.1 | *C.hemisphaerica: Chari et al., 2021 &* *N. vectensis:* Babonis et al., 2022 |
| | | | | Ncol3: STRG.26600 / XP_027042144.1 | *C.hemisphaerica: Denker et al., 2008* & *N. vectensis:* Sunagar et al., 2018 |
| | | | | Nematogalectin-like: STRG.22007 / XP_027037748.1 | *N. vectensis:* Babonis & Martindale, 2017 |

**Supplementary Figure 11: Diagnostic marker genes used for assigning cell type identities in the coral planula atlases.**

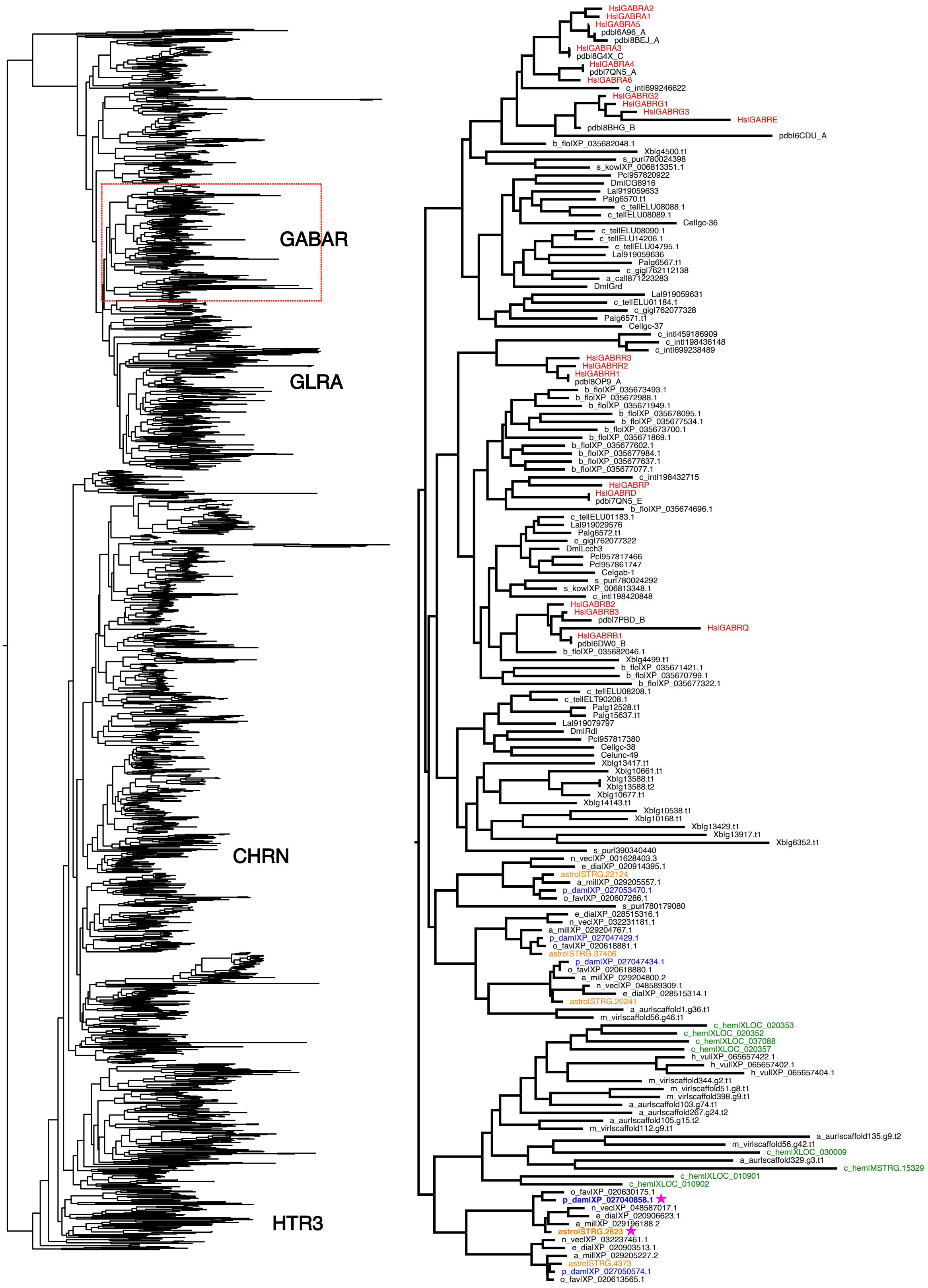
GABAR
GLRA
CHRN
HTR3
HsIGABRA2
HsIGABRA1
HsIGABRA5
pdbI6A96_A
pdbI8BEJ_A
HsIGABRA3
pdbI8G4X_C
HsIGABRA4
pdbI7QN5_A
HsIGABRA6
c_intI699246622
HsIGABRG2
HsIGABRG1
HsIGABRG3
HsIGABRE
pdbI8BHG_B
pdbI6CDU_A
b_floIXP_035682048.1
XbIg4500.t1
s_purI780024398
s_kowIXP_006813351.1
PcI957820922
DmICG8916
LaI919059633
PaIg6570.t1
c_telIELU08088.1
c_telIELU08089.1
CeIIgc-36
c_telIELU08090.1
c_telIELU14206.1
c_telIELU04795.1
LaI919059636
PaIg6567.t1
c_gigI762112138
a_calI871223283
DmIGrd
LaI919059631
c_telIELU01184.1
c_gigI762077328
PaIg6571.t1
CeIIgc-37
c_intI459186909
c_intI198436148
c_intI699238489
HsIGABRR3
HsIGABRR2
HsIGABRR1
pdbI8OP9_A
b_floIXP_035673493.1
b_floIXP_035672988.1
b_floIXP_035671949.1
b_floIXP_035678095.1
b_floIXP_035677534.1
b_floIXP_035673700.1
b_floIXP_035671869.1
b_floIXP_035677602.1
b_floIXP_035677984.1
b_floIXP_035677637.1
b_floIXP_035677077.1
c_intI198432715
HsIGABRP
HsIGABRD
pdbI7QN5_E
b_floIXP_035674696.1
c_telIELU01183.1
LaI919029576
PaIg6572.t1
c_gigI762077322
DmILcch3
PcI957817466
PcI957861747
CeIgab-1
s_purI780024292
s_kowIXP_006813348.1
c_intI198420848
HsIGABRB2
HsIGABRB3
pdbI7PBD_B
HsIGABRQ
HsIGABRB1
pdbI6DW0_B
b_floIXP_035682046.1
XbIg4499.t1
b_floIXP_035671421.1
b_floIXP_035670799.1
b_floIXP_035677322.1
c_telIELU08208.1
c_telIELT90208.1
PaIg12528.t1
PaIg15637.t1
LaI919079797
DmIRdl
PcI957817380
CeIIgc-38
CeIunc-49
XbIg13417.t1
XbIg10661.t1
XbIg13588.t1
XbIg13588.t2
XbIg10677.t1
XbIg14143.t1
XbIg10538.t1
XbIg10168.t1
XbIg13429.t1
XbIg13917.t1
XbIg6352.t1
s_purI390340440
n_vecIXP_001628403.3
e_diaIXP_020914395.1
astroISTRG.22124
a_milIXP_029205557.1
p_damIXP_027053470.1
o_favIXP_020607286.1
s_purI780179080
e_diaIXP_028515316.1
n_vecIXP_032231181.1
a_milIXP_029204767.1
p_damIXP_027047429.1
o_favIXP_020618881.1
astroISTRG.37406
p_damIXP_027047434.1
o_favIXP_020618880.1
a_milIXP_029204800.2
n_vecIXP_048589309.1
e_diaIXP_028515314.1
astroISTRG.20241
a_aurIscaffold1.g36.t1
m_virIscaffold56.g46.t1
c_hemIXLOC_020353
c_hemIXLOC_020352
c_hemIXLOC_037088
c_hemIXLOC_020357
h_vulIXP_065657422.1
h_vulIXP_065657402.1
h_vulIXP_065657404.1
m_virIscaffold344.g2.t1
m_virIscaffold51.g8.t1
m_virIscaffold398.g9.t1
a_aurIscaffold103.g74.t1
a_aurIscaffold267.g24.t2
a_aurIscaffold105.g15.t2
m_virIscaffold112.g9.t1
a_aurIscaffold135.g9.t2
m_virIscaffold56.g42.t1
c_hemIXLOC_030009
a_aurIscaffold329.g3.t1
c_hemIMSTRG.15329
c_hemIXLOC_010901
c_hemIXLOC_010902
o_favIXP_020630175.1
p_damIXP_027040858.1
n_vecIXP_048587017.1
e_diaIXP_020906623.1
a_milIXP_029196188.2
astroISTRG.2823
n_vecIXP_032237461.1
e_diaIXP_020903513.1
a_milIXP_029205227.2
astroISTRG.4373
p_damIXP_027050574.1
o_favIXP_020613565.1

**Supplementary Figure 12: Maximum Likelihood phylogenetic analysis of ligand-gated ion channels.**

The tree on the left includes putative ligand-gated ion channel sequences categorized by including the Pfam domains Neur_chan_LBD and Neur_chan_memb. GABAR: ionotropic GABA receptors; GLRA: Glycine receptors; CHRN = Acetylcholine receptors; HTR3: 5-hydroxytryptamine receptors 3. The tree on the right is a zoomed view of the region outlined by the dotted red box. Sequences from *Clytia*, *Astroides*, and *Pocillopora* are highlighted in green, orange, and blue, respectively, and human sequences are marked in red. Magenta stars indicate expression in GLWamide cells. Hs = *Homo sapiens*; pdb = Protein Data Bank (3D structures); c_int = *Ciona intestinalis*; b_flo = *Branchiostoma floridae*; s_pur = *Strongylocentrotus purpuratus*; s_kow = *Sa*ccoglossus kowalevskii; Pc = *Priapulus caudatus*; Dm = *Drosophila melanogaster*; La = *Lingula anatina*; Pa = *Phoronis australis*; c_tel = *Capitella teleta*; Ce = *Caenorhabditis elegans*; c_gig = *Crassostrea gigas*; a_cal = *Aplysia californica*; Xb = *Xenoturbella bockii*; n_vec = *Nem*atostella *vectensis*; e_dia = *Exaiptasia diaphana*; astro = *Astroides calycularis*; a_mil = *Acropora millep*ora; p_dam = *Pocillopora damicornis*; o_fav = *Orbicella faveolata*; a_aur = *Aurelia aurita*; m_vir = *Morbakka virulenta*; c_hem = *Clytia hemisphaerica*; h_vul = *Hydra vulgaris*.

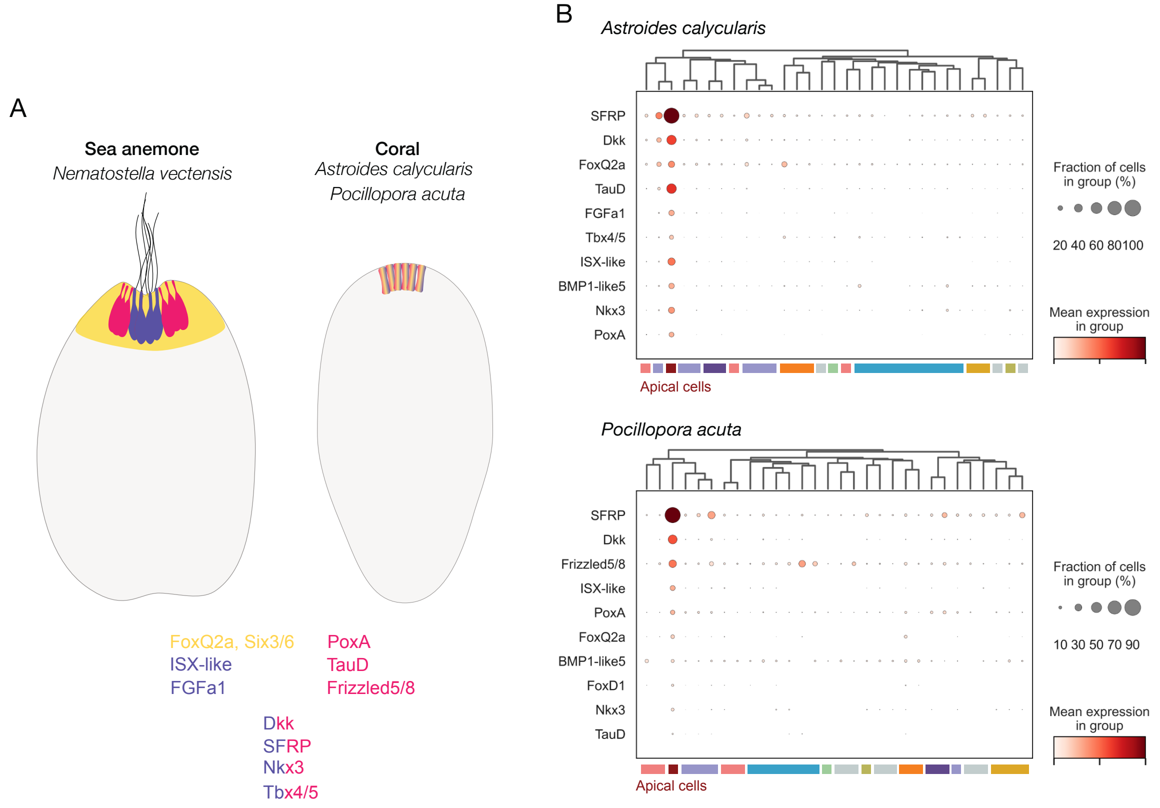


**Supplementary Figure 13: Homology of coral planula apical cells to the apical organ of *Nematostella* planula.**

(A) Schematics illustrating spatial expression of known *Nematostella* apical organ genes. Gene names are color-coded according to their expression in specific areas/cell types. Yellow indicates the apical domain area; purple cells correspond to the apical tuft cells and magenta cells correspond to the surrounding larval neurons. Gene names colored half purple and half magenta indicate expression that is not specifically confined to either of the two apical organ cell types and is presumably present in both. References for spatial gene expression data: FoxQ2a, Six3/6 (Sinigaglia et al., 2013); FGFa1 (Gilbert et al., 2022; Rentzsch et al., 2008; Sinigaglia et al., 2013); ISX-like, PoxA (Gilbert et al., 2022); TauD, Frizzled 5/8, SFRP (Sinigaglia et al., 2015); Dkk (Lee et al., 2006); Nkx3 (Gilbert et al., 2024; Marlow et al., 2013); Tbx4/5 (Gilbert et al., 2024). (B) Dot plots showing expression of the *Nematostella* apical organ markers across *Astroides* (top) and *Pocillopora* (bottom) cell clusters. Gene names are listed on the left and cluster identities are indicated in colored panels at the bottom. The color coding of the clusters corresponds to the cell class annotations in Fig. 4. The color gradient indicates average expression, and the size of the dots indicates the percentage of cells expressing the gene.

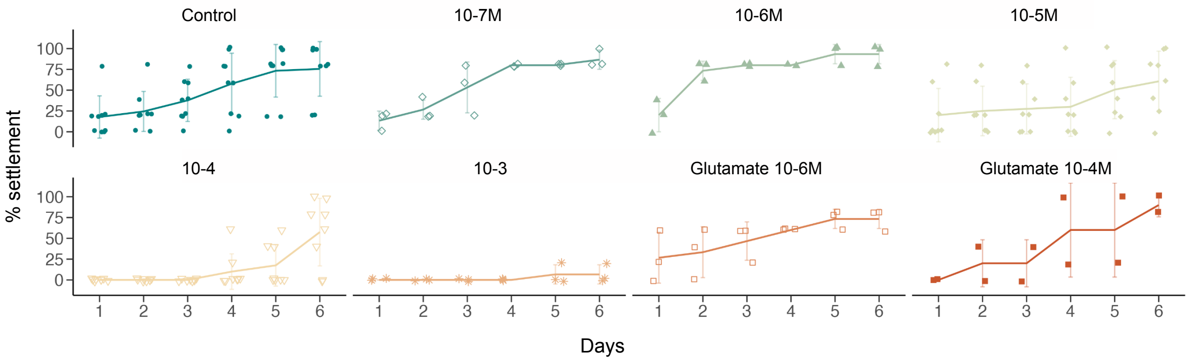


**Supplementary Figure 14 : Effect of Taurine and Glutamate treatment on *Astroides* settlement over time.**

Plots show the percentage of larvae that settled over the course of treatment (days) for each condition. Each dot represents the percentage of larvae that settled in an individual well, with 5 planulae used per treatment condition. Error bars indicate the standard deviation (see raw treatment data in Supplementary table 4).

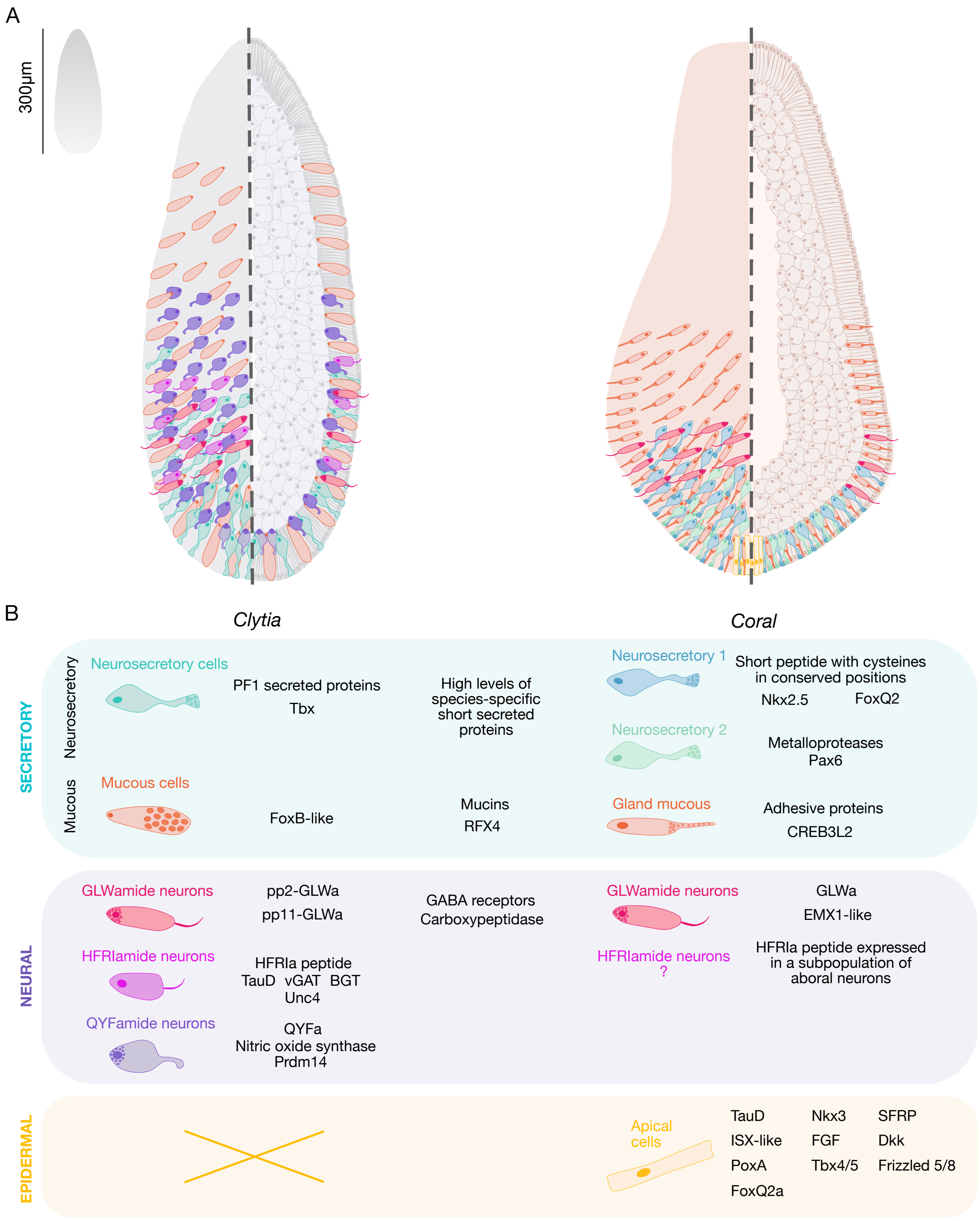


**Supplementary Figure 15: Comparison of specialized aboral cell types between *Clytia* and coral planulae.**

(A) Schematics illustrating the distribution of aboral cell types in *Clytia* and coral planulae. The left section shows a surface view, and the right section corresponds to a medial view. The schematics of the two planulae are depicted at the same size for comparison purposes;

however, the *Clytia* planula is significantly smaller than the *Astroides* planula. (B) Comparison of the molecular signatures of aboral cell types between *Clytia* and coral planulae.

### Supplementary references


Gilbert, E., Craggs, J., & Modepalli, V. (2024). Gene Regulatory Network that Shaped the Evolution of Larval Apical Organ in Cnidaria. *Molecular Biology and Evolution*, *41*(1).

Gilbert, E., Teeling, C., Lebedeva, T., Pedersen, S., Chrismas, N., Genikhovich, G., & Modepalli, V. (2022). Molecular and cellular architecture of the larval sensory organ in the cnidarian Nematostella vectensis. *Development* , *149*(16).

Larkin, M. A., Blackshields, G., Brown, N. P., Chenna, R., McGettigan, P. A., McWilliam, H., Valentin, F., Wallace, I. M., Wilm, A., Lopez, R., Thompson, J. D., Gibson, T. J., & Higgins, D. G. (2007). Clustal W and Clustal X version 2.0. *Bioinformatics (Oxford, England)*, *23*(21), 2947–2948.

Layden, M. J., Boekhout, M., & Martindale, M. Q. (2012). Nematostella vectensis achaete-scute homolog NvashA regulates embryonic ectodermal neurogenesis and represents an ancient component of the metazoan neural specification pathway. *Development* , *139*(5), 1013–1022.

Lee, P. N., Pang, K., Matus, D. Q., & Martindale, M. Q. (2006). A WNT of things to come: evolution of Wnt signaling and polarity in cnidarians. *Seminars in Cell & Developmental Biology*, *17*(2), 157–167.

Levy, S., Elek, A., Grau-Bové, X., Menéndez-Bravo, S., Iglesias, M., Tanay, A., Mass, T., & Sebé-Pedrós, A. (2021). A stony coral cell atlas illuminates the molecular and cellular basis of coral symbiosis, calcification, and immunity. *Cell*, *184*(11), 2973–2987.e18.

Marlow, H., Matus, D. Q., & Martindale, M. Q. (2013). Ectopic activation of the canonical wnt signaling pathway affects ectodermal patterning along the primary axis during larval development in the anthozoan Nematostella vectensis. *Developmental Biology*, *380*(2), 324–334.

Rentzsch, F., Fritzenwanker, J. H., Scholz, C. B., & Technau, U. (2008). FGF signalling controls formation of the apical sensory organ in the cnidarian Nematostella vectensis. *Development* , *135*(10), 1761–1769.

Sabin, K. Z., Chen, S., Hill, E. M., Weaver, K. J., Yonke, J., Kirkman, M., Redwine, W. B., Klompen, A. M. L., Zhao, X., Guo, F., McKinney, M. C., Dewey, J. L., & Gibson, M. C.

(2024). Graded FGF activity patterns distinct cell types within the apical sensory organ of the sea anemone Nematostella vectensis. *Developmental Biology*, *510*, 50–65.

Sinigaglia, C., Busengdal, H., Leclère, L., Technau, U., & Rentzsch, F. (2013). The bilaterian head patterning gene six3/6 controls aboral domain development in a cnidarian. *PLoS Biology*, *11*(2), e1001488.

Sinigaglia, C., Busengdal, H., Lerner, A., Oliveri, P., & Rentzsch, F. (2015). Molecular characterization of the apical organ of the anthozoan Nematostella vectensis. *Developmental Biology*, *398*(1), 120–133.

Steger, J., Cole, A. G., Denner, A., Lebedeva, T., Genikhovich, G., Ries, A., Reischl, R., Taudes, E., Lassnig, M., & Technau, U. (2022). Single-cell transcriptomics identifies conserved regulators of neuroglandular lineages. *Cell Reports*, *40*(12), 111370.
